# Humanizing a CD28 signaling domain affects CD8 activation, exhaustion and stem-like precursors

**DOI:** 10.1101/2025.03.10.642460

**Authors:** Alexander E. Brady, Shankar Revu, Dongwen Wu, Hannah Fisk, Khadija Kone, Alexandria Lydecker, Elliott J. Purser, Norah Smith, Zachary T. Hilt, Sarah Woodyear, Sarah Caddy, Sebastien Gingras, Brian Rudd, Mandy M. McGeachy

## Abstract

CD28 ligation provides critical signals that modulate activated T cell fate. In a human to mouse reverse-engineering approach, a single amino acid substitution adjacent to the C-terminal proline-rich domain created CD28^A210P^ mice with enhanced signaling. CD28^A210P^ mice experienced pro-inflammatory responses to CD28 superagonist antibody, analogous to severe cytokine storm induced in a human clinical trial, with a striking increase of activated CD8 T cells. In acute and chronic viral infections, early activation and expansion of CD28^A210P^ CD8 effector T cells increased, with accelerated exhaustion in chronic infection. Mechanistically, CD28^A210P^ enhanced JunB, IL-2, and inhibitory receptors driven by MEK1/2. Generation of CD28^A210P^ stem-like progenitor (Tpex) cells was enhanced in acute and chronic infections, and further expanded by PD-L1 blockade in chronically-infected mice. Thus, ‘humanized’ PYAP mice reveal key roles for CD28 signaling strength in CD8 activation, accelerating exhaustion during antigen persistence, while promoting and sustaining Tpex during acute and chronic viral infection.

**One sentence Summary:** A single amino acid substitution adjacent to PYAP to ‘humanize’ CD28 signaling enhances superagonist response, early CD8 activation and Tpex generation during viral infection while accelerating exhaustion and sustaining Tpex during chronic infection.

Graphical Abstract:
‘Humanized’ CD28 PYAPP enhances numbers of CD8 T cell effectors and stem-like precursors during acute viral infection, and accelerates exhaustion while sustaining increased self-renewing Tpex cells that are favored during PD-L1 blockade.

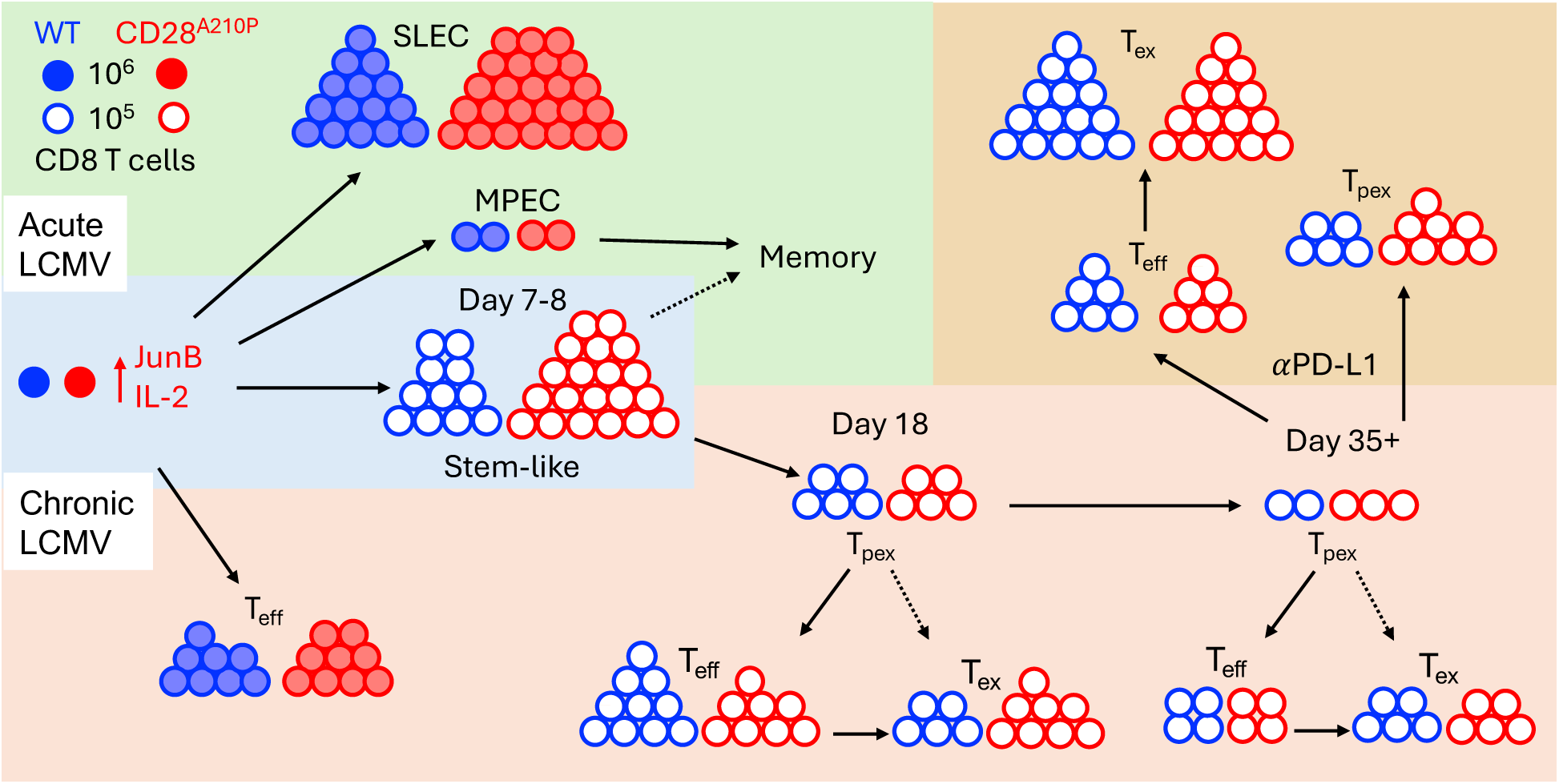

## Introduction

CD28 is a constitutively expressed T cell costimulatory receptor critical for co-activating TCR signaling to facilitate functional changes including IL-2 production, metabolic shifts, proliferation, and survival. It has become increasingly clear that nuanced CD28 signaling dynamically alters T cell fate throughout an immune response, rather than providing a simple ‘go/no-go’ signal. The cytoplasmic tail of CD28 contains three well-characterized signaling domains that are highly conserved amongst mammals (*1–5*). The relative contributions and exact signaling downstream of each domain have been challenging to decipher, in part because of the shared interdependent circuitry of TCR and CD28 signaling molecules. Differences in experimental systems used to study CD28 function have further complicated interpretation. For example, early experiments in the human T cell Jurkat cell line may have over-attributed the role of the proximal PI3K-binding domain and Akt in CD28 signaling due to Jurkat’s lack of critical negative regulators PTEN and SHP1 (*5*). Results in mice bearing targeted loss-of-function mutations indicated greater importance of the C-terminal proline-rich PYAP domain in functional outcomes of CD28 signaling on T cell activation in vitro and in vivo. The defects in thymic development of mature T cells caused by loss-of-function and global knockouts of CD28, including FoxP3^+^ regulatory T cells, create additional complications for understanding CD28 function in the periphery (*6–8*). However, conditional deletion of CD28 in mature T cells confirmed the continued requirement for CD28 signaling in the antigen-experienced T cell response to checkpoint blockade and in sustaining stem-like T cells during chronic antigen stimulation(*9, 10*) (*11, 12*).

The central role of CD28 signaling in T cells makes it an attractive target for immunotherapeutics. CTLA-4Ig competes for the CD28 ligand B7 to reduce T cell activation and is beneficial in autoimmune diseases like Rheumatoid Arthritis (*13*); direct antagonism of CD28 is also under investigation for transplantation tolerance (*14*). On the other hand, monoclonal antibodies against the inhibitory receptors PD-1 and CTLA4 are being used to enhance TCR/CD28 signaling for cancer therapy (*15*). CD28 signaling domains have been incorporated into chimeric antigen receptors (CAR) for CAR-T therapies that are revolutionizing cancer therapy and are under investigation for other chronic diseases such as multiple sclerosis, lupus, and HIV/AIDS (*16–18*). Although not without side effects, these FDA approved therapies have provided life changing clinical benefit for patients and continue to be refined and expanded on for new applications in the clinic.

In contrast to CD28-based therapeutic successes, an attempt to use CD28 autonomous activation with CD28-superagonist (CD28SA) for tolerance induction became a dramatic example of interspecies differences that alter immunity. Pre-clinical mouse studies suggested CD28SA had potential to induce immunoregulation in settings of autoimmunity and allotransplantation by inducing significant expansion of Tregs (*19, 20*) (*21*). Safety studies in rhesus macaques provided support to proceed to phase I clinical trials in healthy young adult male volunteers. Shockingly, all participants receiving CD28SA drug experienced rapid and severe cytokine storm requiring hospitalization and life-saving interventions (*22*). Since then, several factors have been proposed to explain this unexpected outcome. Mice used for preclinical studies were housed in ultraclean specific-pathogen free facilities typical for scientific research settings, and we now know that compared to humans who experience a diverse microbial environment, these mice have an immature immune system with barely any tissue-circulating or resident T cells (*23*). ‘Wildling’ lab mice born to wild-caught mothers through embryo transfer confirm that CD28SA drives less Tregs and more weight loss in animals with a diverse microbiota, though these mice did not experience the severity of cytokine storm induced in human volunteers (*24*). In addition, rhesus macaques were found to downregulate CD28 on memory cells to a greater degree than humans, which could have prevented activation by CD28SA (*25–27*).

Interspecies differences in the CD28 molecule itself may have also contributed to the disastrous clinical trial of CD28SA. Although the PYAP domain is highly conserved, the adjacent amino acid varies: mice have an alanine (PYAP<u>A</u>), humans have an additional proline (PYAP<u>P</u>). The prolines are critical to the PYAP domain function, and AYAA mice have severe impairments in T cell development and activation (*7, 28, 29*). In a ground-breaking study, Porciello et al showed that simply substituting the amino acid and expressing CD28-PYAPA in human CD4 T cells or CD28-PYAPP in mouse CD4 T cells was sufficient to alter signaling and cytokine response to CD28SA in vitro (*30*).

To probe the outcomes of enhanced CD28 on T cell function in vivo, we generated CD28^A210P^ mice with a ‘humanized’ PYAPP domain. In contrast to CD28 deletion and loss of function approaches, enhancing CD28 signaling did not alter thymic development or Treg numbers. Challenging CD28^A210P^ mice with CD28SA drove a pro-inflammatory response alongside a striking increase in CD8 T cell activation previously underappreciated from mouse studies of CD28SA. We therefore sought to determine how enhanced CD28 signaling influences more physiological responses to acute and chronic viral infection. Here we report that enhancing CD28 signaling increases the numbers of early effector and stem-like progenitor (Tpex) CD8 T cells, and in the case of chronic infection accelerated CD8 T cell exhaustion. Mechanistically, CD28^A210P^ T cells have enhanced induction of JunB and required MEK1/2 for increased activation and early upregulation of inhibitory co-receptors. During late-stage chronic infection, CD28^A210P^ Tpex were sustained at higher numbers while effector/exhausted cells were similar to WT. PD-L1 blockade revealed that enhanced CD28 signaling better supports Tpex but did not provide an advantage for driving effectors. Together, these data support a model in which CD28 co-stimulation is critical for early numbers of effector and stem-like progenitor cells that may later require additional intervention to rejuvenate chronically stimulated T cells.

## Results

### Generation of a CD28^A210P^ ‘humanized’ mouse

The CD28 PYAP domain is highly conserved amongst four-limbed vertebrates, and we found that the adjacent C-terminal amino acid has diverged over time, with Eutherian mammals with alanine at that position (PYAPA) while marsupials appear to retain threonine (PYAPT) similar to birds and crocodiles (**Fig 1A**). Primates have undergone a further substitution to proline (PYAPP) (**Fig 1A**), previously shown to enhance CD28 autonomous signaling in human versus mouse CD4 T cells (*30*). To test the role of enhanced CD28 signaling in murine T cell models, we generated CD28^A210P^mice with a proline to alanine substitution at position 210 of CD28, resulting in a PYAPP domain as found in humans (**Fig 1B**, S1A). CD28^A210P^ mice breed normally, display similar body weights to WT mice (Fig S1Q), and do not display overt signs of inflammatory disease or other health issues. Given the requirement for CD28 PYAP signaling in thymic T cell development, we asked if enhancing CD28 signaling via the CD28 A210P mutation alters T cell maturation and/or subsequent seeding of secondary lymphoid organs. CD28^A210P^ mice displayed normal thymic T cell development characterized by similar frequencies and numbers of double negative (1–4), double positive, CD4 single positive, CD8 single positive, and Foxp3^+^ Treg populations (**Fig 1C, 1D**, S1B-E) compared to WT mice. As T cells undergo positive selection CD69 and TCR-β are upregulated and as CD4SP and CD8SP T cells are selected, CD69 is subsequently downregulated, a process altered in B7 dKO mice (*7*). Enhancing CD28 signaling does not alter frequencies or numbers of T cells in each stage of development (Fig S1F). CD5 GMFI across thymocyte populations in CD28^A210P^ mice is similar to WT indicating equivalent strength of TCR signaling in developing thymocytes (**Fig 1E**). Functional CD28:B7 is required for thymic clonal deletion (*7*), however there was no alteration in CD28^A210P^ thymic clonal deletion or death by neglect compared to WT (Fig S1G). Correlating to normal thymic development, secondary lymphoid organs in CD28^A210P^ mice also display normal homeostatic T cell populations as characterized by frequencies and numbers of naïve, effector memory, and central memory CD4^+^ and CD8^+^ T cells compared to WT mice (**Fig 1F**, S1H-O). Treg (FOXP3^+^ CD4^+^) cell frequencies and numbers are also normal in spleens and lymph nodes of CD28^A210P^ mice compared to WT (**Fig 1G**, S1P). We confirmed the A210P mutation does not alter CD28 surface expression on homeostatic T cell subsets (**Fig 1H**). Together with published data on CD28 loss of function mutants, these data suggest that CD28 signaling is required for normal T cell development, but enhancing C-terminal signaling capacity of CD28 does not overtly change homeostatic T cell output.

**Figure 1).**
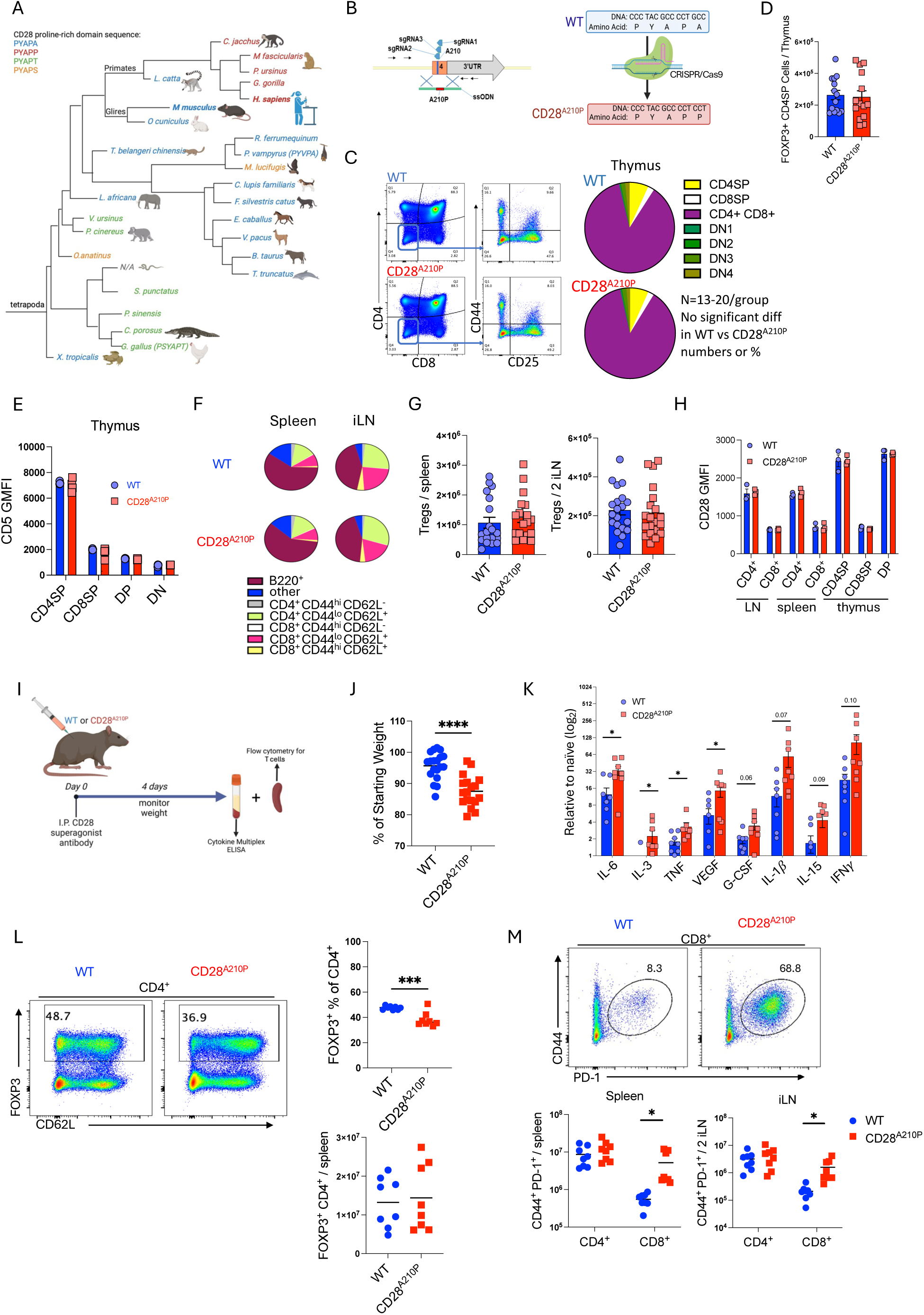
Generation of a CD28^A210P^ ‘humanized’ mouse that better recapitulates the human response to CD28 superagonist antibody. **A)** Phylogenetic tree of CD28 C-terminal proline-rich domain amino acid sequence in tetrapod mammals, generated from uniprot sequence database(*62*). **B)** CD28^A210P^ mice were generated by CRISPR-mediated substitution of proline for alanine at amino acid position 210 of mouse CD28. Gene and amino acid sequence of WT and CD28^A210P^ CD28. **C-E**: Thymus of adult CD28^A210P^ and WT mice assessed by flow cytometry for T cell development stage subsets (C), thymic FoxP3^+^ Tregs (D) and surface CD5 expression (E). **F-H**: Peripheral LN and spleen of adult WT and CD28^A210P^ mice were assessed by flow cytometry for proportions of T cell subsets (F), Foxp3^+^ Tregs (G) and surface expression of CD28 (H). **I-M**: Mice were injected with CD28 superagonist (CD28SA) and monitored for weight loss (J), on day 4 post-injection cytokines were assessed in serum (K), and spleen analyzed by flow cytometry for frequencies and absolute counts of Foxp3^+^ Tregs (L) and activated CD4^+^ and CD8^+^ T cells (M). Data are pooled from at least two experiments with 3-6 per group, except E&H are one experiment representative of two, dots show individual mice, significance assessed by Student’s t-test or one-way ANOVA, * = p < 0.05, ** = p < 0.01, *** = p < 0.001.

### CD28^A210P^ mice undergo pro-inflammatory response to CD28 superagonist antibody

To determine if enhancing CD28 signaling in mice would better recapitulate the human reaction to a CD28 superagonist antibody we injected WT and CD28^A210P^ mice with superagonist anti-CD28 (D665) (**Fig 1I**). Following injection, CD28^A210P^ mice but not WT mice underwent significant weight loss (**Fig 1J**). CD28^A210P^ serum contained increased levels of systemic pro-inflammatory cytokines that are highly elevated during cytokine storm in humans (*31*), such as IL-6 and TNF compared to WT (**Fig 1K**). Following CD28SA injection absolute cell counts of spleens and inguinal lymph nodes were similar in WT and CD28^A210P^ mice (Fig S2A,L), however we observed decreased frequencies but similar numbers of expanded Tregs in CD28^A210P^ mice compared to WT (**Fig 1L**). Coinciding with lower Treg frequencies, CD28^A210P^ mice showed a greater proportion of activated FOXP3^-^ CD4^+^ T cells (Fig S2J). Strikingly, in WT mice, CD28 superagonist induced a relatively small response in CD8^+^ T cells compared to CD4^+^ (**Fig 1M**). However, CD28^A210P^ activated CD8^+^ T cells were significantly increased in superagonist-treated CD28^A210P^ mice (**Fig 1M**, S2B-I, S2Q-T) suggesting enhancing CD28 signaling via the A210P mutation enables a substantial CD8^+^ T cell response to CD28SA. These data support a predominantly pro-inflammatory response in CD28^A210P^ mice in contrast to the dominant regulatory response seen in WT mice and highlight an unappreciated ability of CD28SA to activate CD8^+^ T cells.

### Enhancing CD28 C-terminal signaling increases initial effector T cell differentiation during acute and chronic viral infection

CD28 superagonist antibodies are unique in that they do not require TCR stimulation (signal 1) to activate T cells. Physiological T cell activation requires both TCR stimulation and CD28 ligation (signal 1 and 2). To examine if enhancing CD28 signaling would also promote effector T cell activation in a more physiological context we utilized LCMV as a model of viral infection and effector T cell activation (**Fig 2A**). 7 days post infection spleens were collected and analyzed by flow cytometry (**Fig 2B**). Enhanced CD28 signaling increased numbers of activated (CD44^+^) (**Fig 2C**) and H-2D^b^ LCMV GP33-specific CD8^+^ T cells (**Fig 2D**, S3A). There were increased CD8^+^ effector T cells expressing CX3CR1 and KLRG1 in CD28^A210P^ spleens compared to WT (**Fig 2E**, S3B). To verify the increased expression activation markers seen in CD28^A210P^ T cells reflects increased effector potential, we performed intracellular cytokine staining on ex vivo LCMV GP33 peptide restimulated splenocytes from WT and CD28^A210P^ mice. Following peptide re-stimulation CD28^A210P^ CD8^+^ T cells showed increased frequencies and numbers of pro-inflammatory cytokine producing cells confirming their greater effector potential (**Fig 2F**, S3C-D). Despite this, CD28^A210P^ mice did not have overt signs of immunopathology or increased weight loss (**Fig 2G**) presumably due to the rapid clearance of virus that occurs with this strain.

**Figure 2).**
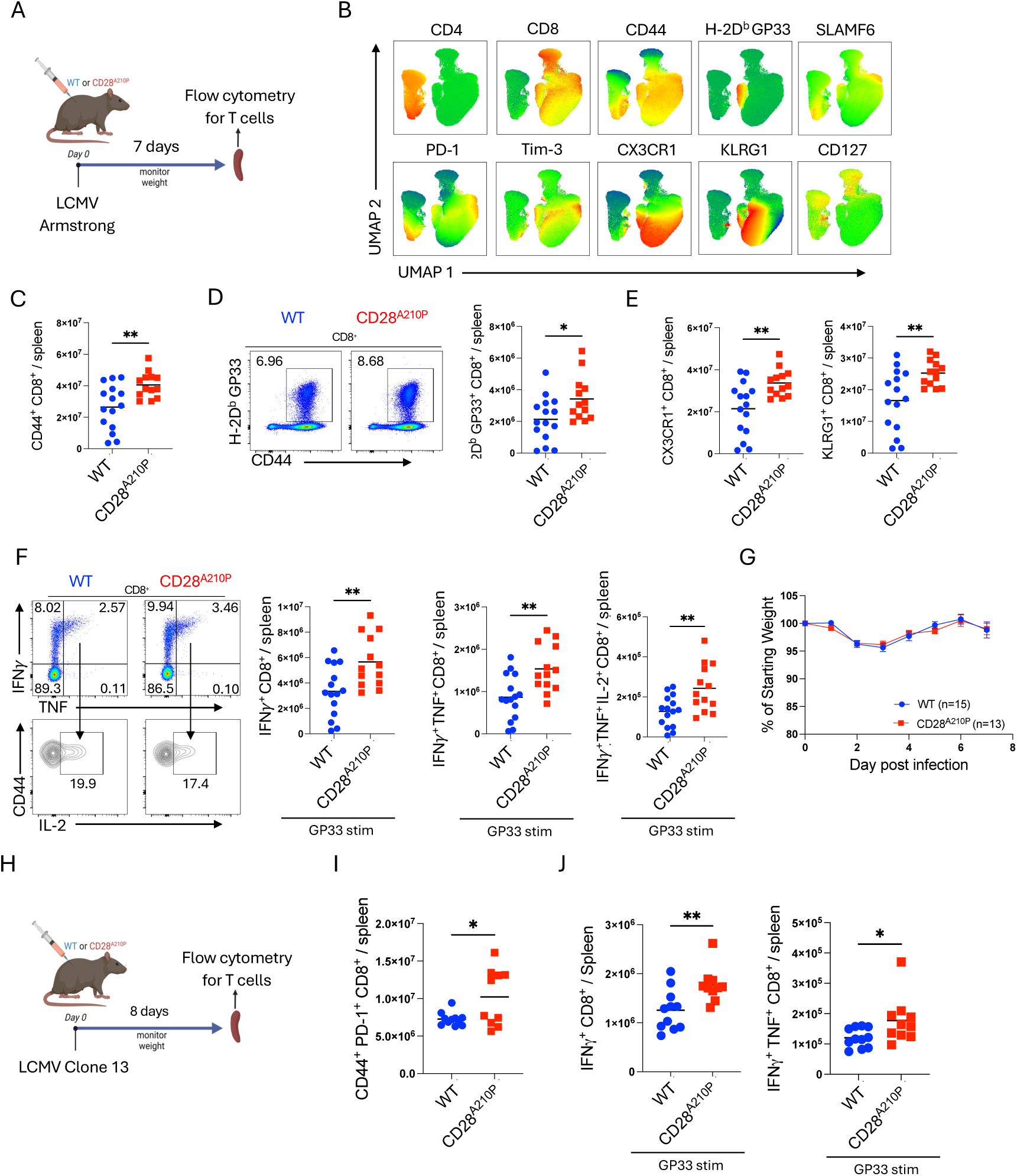
Enhancing CD28 C-terminal signaling increases initial effector T cell differentiation during acute and chronic viral infection. A) Experimental timeline for LCMV Armstrong infection and splenocyte analysis 7 days post infection. B) UMAP clustering and marker heatmap overlay of CD4^+^ and CD8^+^ cells by flow cytometry. Data pooled from WT and CD28^A210P^ splenocytes to construct representative clustering. C) Absolute numbers of CD44^+^ CD8^+^ splenocytes. D) Representative gating and quantification of H-2D^b^ GP33^+^ CD8^+^ T cells. E) Absolute numbers of effector CX3CR1^+^ and KLRG1^+^ CD8^+^ T cells. F) Representative gating and absolute numbers of cytokine producing CD8^+^ T cells following 4-hour ex vivo stimulation with LCMV peptide GP33. G) Weight loss following LCMV Armstrong infection shown as percentage of starting weight on day 0. H) Experimental timeline of LCMV clone 13 infection experiments. I) Absolute numbers of activated (CD44^+^ PD-1^+^) CD8^+^ T cells 8 days post infection with LCMV clone 13. J) Absolute numbers of IFNγ and IFNγ TNF producing CD8^+^ T cells following 4-hour ex vivo stimulation with LCMV peptide GP33. Data are pooled from at least two experiments with 4-8 per group. Significance assessed by Student’s t-test or one-way ANOVA, * = p < 0.05, ** = p < 0.01, *** = p < 0.001.

We then asked if enhancing CD28 signaling during chronic infection would also result in increased effector function. At 8 days post infection with the chronic LCMV clone 13 strain (**Fig 2H**), CD28^A210P^ CD8^+^ T cells showed increased activation (CD44^+^ PD-1^+^) (**Fig 2I**) and IFNγ– and IFNγ/TNF-production potential following peptide re-stimulation, indicating enhanced peak effector differentiation compared to WT (**Fig 2J**, S3G-I). These data together show enhancing CD28 signaling increases initial CD8^+^ T cell effector differentiation during viral infection.

### Enhancing CD28 C-terminal signaling induces early upregulation of CD8^+^ T cell inhibitory receptors and expedites exhaustion

Immunomodulation is required to prevent severe immunopathology during LCMV clone 13 infection (*32*). T cells possess cell-intrinsic means of immunoregulation through upregulation of inhibitory receptors which can be beneficial or detrimental during chronic antigen exposure depending on context (*33*). Following infection with LCMV clone 13 (**Fig 3A**), enhancing CD28 signaling through the CD28 A210èP mutation resulted in increased frequencies and numbers of early PD-1^+^ Tim3^+^ CD8^+^ T cells by day 8 of infection (**Fig 3B**, S3J). Although a majority of these are likely effectors and not bona-fide exhausted cells at this early time point, the increased presence of inhibitory receptors suggests enhanced CD28 signaling promotes strong immunoregulation in parallel to an enhanced effector response (**Fig 2**). Early CD28^A210P^ CD8^+^ PD-1^+^ Tim3^+^ cells displayed higher PD-1 GMFI (**Fig 3C**) indicating enhanced T cell-intrinsic immunoregulation during the effector phase of the T cell response. Coinciding with enhanced immunoregulation at day 8 post LCMV infection, systemic IFNγ is decreased in CD28^A210P^ mice compared to WT mice (**Fig 3D**). Although exhausted cells are low in numbers at this time, CD28^A210P^ T_eff_/T_ex_ ratio is similar to WT (**Fig 3E**). In agreement with increased effector T cell response (**Fig 2**), CD28^A210P^ mice lose significantly more weight than WT during the second week of infection (**Fig 3F**). But in parallel with enhanced inhibitory receptor upregulation and decreased systemic IFNγ, CD28^A210P^ mice quickly recover to weights comparable to WT controls by end of that second week (**Fig 3F**).

**Fig 3).**
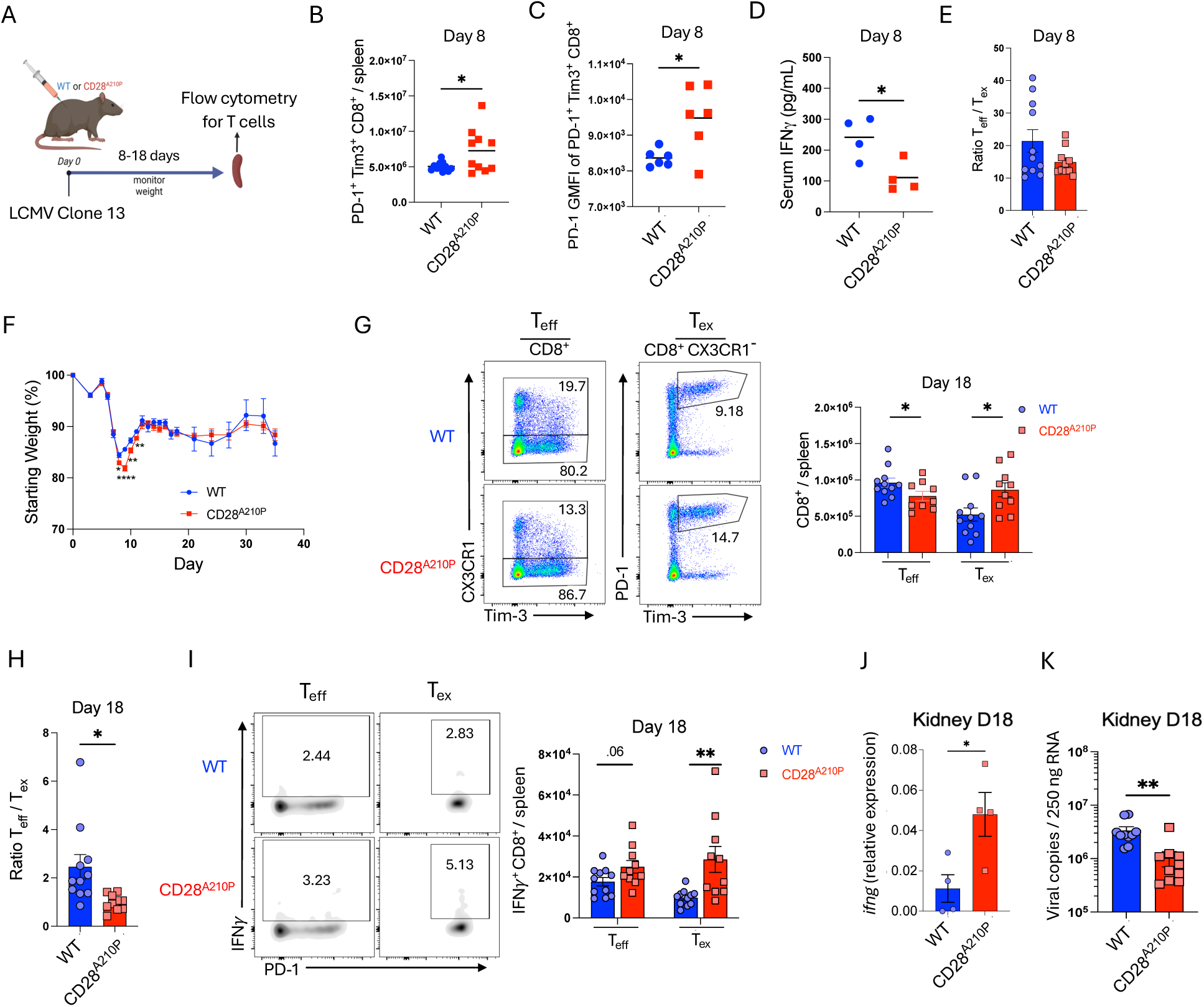
Enhancing CD28 C-terminal signaling induces early upregulation of CD8^+^ T cell inhibitory receptors and expedites exhaustion. **A)** Experimental timeline of LCMV clone 13 infection. **B)** Absolute numbers of splenic PD-1^+^ Tim-3^+^ CD8^+^ T cells 8 days post LCMV clone 13 infection. **C)** PD-1 GMFI of PD-1^+^ Tim-3^+^ CD8^+^ cells 8 days post infection. **D)** Serum IFNγ 8 days post infection. **E)** Ratio of CD8^+^ T_eff_ (CX3CR1^+^) to T_ex_ (CX3CR1^-^ PD-1^+^ Tim-3^+^) splenocytes. **F)** Weight loss following LCMV infection, shown as a percentage of starting weight on day 0, data pooled from multiple experiments, (WT n= 9-28/time point) (CD28^A210P^ n= 8-25/time point) student’s t test used for each day. **G)** Representative gating and quantification of absolute numbers of CD8^+^ effectors (CX3CR1^+^) and exhausted cells (CX3CR1^-^ PD-1^+^ Tim-3^+^) on day 18 post infection. **H)** Ratio of absolute numbers of CD8^+^ CX3CR1^+^ effectors/ CD8^+^ CX3CR1^-^ PD-1^+^ Tim-3^+^ exhausted cells. **I)** Representative gating and quantification of absolute numbers of IFNγ^+^ CD8^+^ T cells of indicated subsets based on previously described surface marker gating. Splenocytes were stimulated ex vivo for 4 hours with LCMV GP33 peptide in the presence of GolgiPlug followed by FACS staining. **J)** RT-qPCR quantification of *ifng* from LCMV clone 13 infected mouse kidney 18 days post infection. **K)** RT-qPCR quantification of viral copies in Kidney tissue from LCMV clone 13 infected mice 18 days post infection. Data are pooled from at least two experiments with 3-6 per group, except C and D one experiment representative of two and J is one experiment. Dots show individual mice. Significance assessed by Student’s t-test or one-way ANOVA, * = p < 0.05, ** = p < 0.01, *** = p < 0.001.

To examine how enhancing CD28 signaling would influence CD8^+^ T cell effector output and exhaustion at an intermediate point of chronic infection, we utilized LCMV clone 13 infection and analyzed effector and exhausted T cells 18 days post infection. Although enhanced CD28 signaling induces a greater effector CD8^+^ response 8 days post infection (**Fig 2**), by day 18 post infection CD28^A210P^ mice display less effectors (CX3CR1^+^) and more exhausted (CX3CR1^-^ PD-1^+^ Tim-3^+^) CD8^+^ cells compared to WT mice (**Fig 3G**, S4A), which results in a decreased effector: exhausted CD8^+^ cell ratio (**Fig 3H**, S4B). Unexpectedly, upon ex vivo GP33 peptide restimulation, CD28^A210P^ CD8^+^ T cells displayed increased IFNγ production potential (**Fig 3I**, S4C), despite skewing towards exhaustion surface markers (**Fig 3G**). Kidney tissue from WT and CD28^A210P^ mice showed enhanced CD28 signaling resulted in increased expression of *ifng* and better viral control (**Fig 3J,K**). These data together suggest although enhancing CD28 signaling increases the initial CD8^+^ effector response, it also drives increased early self-regulation through inhibitory receptors.

### The CD28 A210P mutation enhances MEK1/2-dependent induction of JunB, IL-2, and PD-1/Tim-3 following T cell activation

To verify enhanced the CD28 A210P mutation increased T cell activation in a T-cell intrinsic manner, we assessed IL-2 production as a direct outcome of CD28 signaling. Upon in vitro stimulation with plate-bound agonistic anti-CD3 and anti-CD28 antibodies, primary CD8^+^ T cells from CD28^A210P^ mice produced more IL-2 than WT CD8^+^ T cells (**Fig 4A**). Following LCMV infection CD28^A210P^ CD8^+^ T cells displayed increased inhibitory receptor expression (**Fig 3**). To assess if enhanced CD28-induced inhibitory receptor upregulation is CD8^+^ T cell-intrinsic we activated WT and CD28^A210P^ CD8^+^ T cells with αCD3/αCD28 and assessed PD-1 and Tim-3. Following activation with αCD3/αCD28 together, but not αCD3 alone, CD28^A210P^ CD8^+^ T cells displayed increased frequencies of PD-1^+^ Tim-3^+^ cells (**Fig 4B**), confirming the role of enhanced CD28 in induction of inhibitory receptors. To determine how the CD28 A210P mutation alters signaling downstream of CD28 we assessed signaling molecules and transcription factors known to be downstream of CD28. There were no differences between WT and CD28^A210P^ cells in p-Akt, p-S6, and IκBα protein regulation following T cell activation (Fig S5A-C). We then probed AP-1 family members, which have been shown to be critical for early CD28-dependent T cell activation and dynamically regulate T cell responses in chronic antigen settings (*34–36*). Following 2 hours of αCD3/αCD28 stimulation, but not αCD3 alone, CD28^A210P^ CD8^+^ T cells displayed increased nuclear abundance of the AP-1 family member JunB (**Fig 4C**), but not IRF4 or BATF (Fig S5D-E).

**Figure 4).**
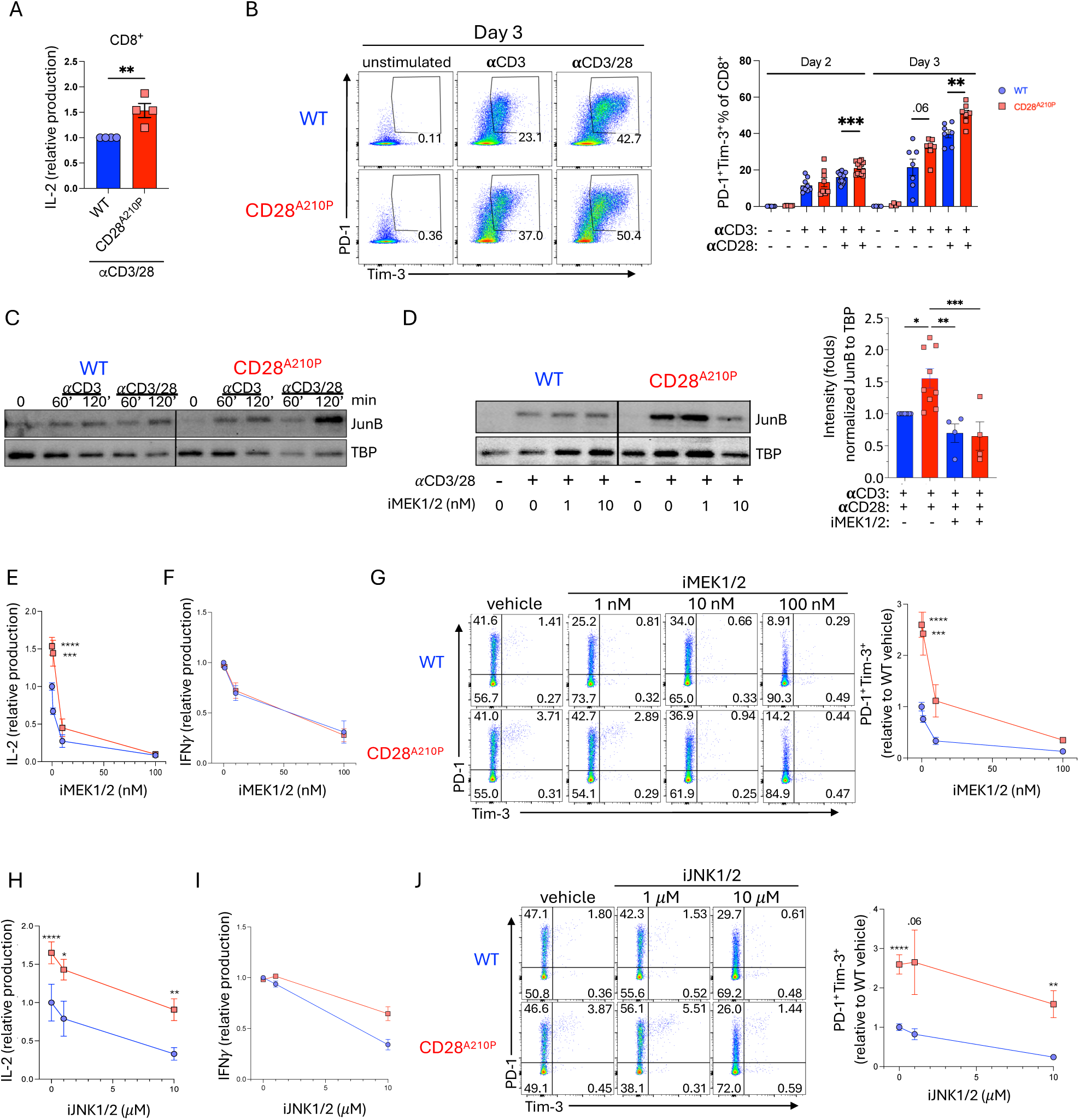
The CD28 A^210^P mutation enhances MEK-dependent induction of JunB, IL-2, and PD-1/Tim-3 following T cell activation. **A-J)** CD8^+^ T cells cells were stimulated with plate-bound anti-CD3 and anti-CD28 agonistic antibodies (1μg/mL in A,B E-J) (5μg/mL in C,D). **A)** Supernatant IL-2 was quantified by ELISA following 6 hours of stimulation. Each point represents the mean of replicate wells from an independent experiment. Data normalized to WT control in each independent experiment. **B)** Representative gating and quantification of PD-1 and Tim-3 on CD8^+^ cells stimulated for 2-3 days. Day 2: points indicate replicate wells pooled from 3 independent experiments. Day 3: points indicate replicate wells pooled from 2 independent experiments. **C)** Representative immunoblot of JunB from WT or CD28^A210P^ CD8^+^ T cell nuclear extracts following stimulation for indicated times. Representative of 5 independent experiments. **D)** Representative immunoblot of JunB from nuclear extracts from WT or CD28^A210P^ CD8^+^ T cells stimulated 2 hours in the presence of DMSO vehicle control or indicated concentrations of the MEK1/2 inhibitor, trametinib. Representative of 3 independent experiments. Quantification of vehicle treated extracts is pooled from 9 independent experiments and inhibitor treated extracts are pooled from 4 independent experiments. **E)** CD8^+^ cells were stimulated for 6 hours in the presence of vehicle or indicated concentrations of the MEK1/2 inhibitor. Supernatant IL-2 was quantified by ELISA. Data normalized to WT control in each independent experiment. Points indicate mean of pooled experimental replicates from each experiment. **F)** CD8^+^ cells were stimulated for 24 hours. Vehicle or MEK1/2 inhibitor were added to wells prior to cell seeding. Supernatant IFNγ was quantified by ELISA. Data normalized to WT control in each independent experiment. Points indicate mean of pooled experimental replicates from each experiment. **G)** Representative gating and quantification of PD-1 and Tim-3 on CD8^+^ cells stimulated for 24 hours with vehicle or MEK1/2 inhibitor added to wells prior to cell seeding. Data normalized to WT control in each independent experiment. Points indicate mean of pooled replicates from 2 independent experiments. **H)** CD8^+^ T cells were stimulated for 6 hours. Vehicle or indicated concentrations of the JNK1/2 inhibitor, JNK-IN-8, were added to wells prior to cell seeding. Supernatant IL-2 was quantified by ELISA. Data normalized to WT control in each independent experiment. Points indicate mean of pooled experimental replicates from 3 independent experiments. **I)** CD8^+^ cells were stimulated for 24 hours. Vehicle or JNK1/2 inhibitor were added to wells prior to cell seeding. Supernatant IFNγ quantified by ELISA. Data normalized to WT control in each independent experiment. Points indicate mean of pooled experimental replicates from 3 independent experiments. **J)** Representative gating and quantification of PD-1 and Tim-3 on CD8^+^ T cells stimulated 24 hours with vehicle or JNK1/2 inhibitor added prior to cell seeding. Data normalized to WT control in each independent experiment. Points indicate mean of pooled replicates from 2 independent experiments. Data are pooled or representative of at least two experiments with significance assessed by Student’s t-test or one-way ANOVA, * = p < 0.05, ** = p < 0.01, *** = p < 0.001.

We next sought to determine the upstream signaling leading to enhanced JunB induction in CD28^A210P^ CD8^+^ T cells. CD28-dependent MEK1/2 activity has been implicated in the induction of AP-1/JunB and subsequent IL-2 production in T cells (*37–41*). Our data indicate JunB is increased due to enhanced CD28 C-terminal signaling. We then asked if a low dose of MEK1/2 inhibitor could bring down CD28^A210P^ nuclear JunB to WT levels following αCD3/αCD28 stimulation. Primary WT and CD28^A210P^ CD8^+^ T cells were activated with plate-bound αCD3/αCD28 in the presence of vehicle or low dose of the FDA-approved MEK1/2 inhibitor, trametinib, for 2 hours. MEK1/2 inhibition decreased CD28^A210P^ nuclear JunB to a similar level to that present in WT T cells (**Fig 4D**). Correspondingly, MEK1/2 inhibition normalized IL-2 production by CD28^A210P^ CD8^+^ T cells to a level similar to WT (**Fig 4E**). Interestingly, enhanced CD28 signaling does not increase IFNγ production within the first 24 hours of CD8^+^ T cell activation (**Fig 4F**). Given this result we reasoned that the increased IFNγ^+^ cells seen in CD28^A210P^ mice following LCMV infection (**Fig 2F**) is due to increased IL-2-driven proliferation of effector cells rather than increased IFNγ production per cell.

As observed in vivo, inhibitory receptors were more upregulated in CD28^A210P^ T cells (**Fig 4B**). Heterodimers of cFos and Jun family members, increase expression of inhibitory receptors such as PD-1 and Tim-3 (*42, 43*). We asked if low dose MEK1/2 inhibition could decrease PD-1 and Tim-3 in CD28^A210P^ T cells to similar levels to WT T cells. Following 24 hours of αCD3/αCD28 activation in the presence of 10 nM MEK1/2 inhibitor, CD28^A210P^ T cells displayed similar levels of PD-1 and Tim-3 compared to vehicle treated WT T cells (**Fig 4G**).

The canonical AP-1 heterodimer consisting of cJun and cFos, requires CD28 signaling and has been extensively implicated in T cell activation and effector function and contributes to CD28-dependent IL-2 production (*34, 44*). cJun activity requires phosphorylation by JNK while JunB is less dependent on JNK phosphorylation. To examine if enhanced AP-1 signaling in CD28^A210P^ T cells also invoked enhanced JNK-dependent activity we titrated the JNK inhibitor, JNK-IN-8, into CD8^+^ T cell activation experiments and assessed IL-2 and IFNγ production and upregulation of PD-1 and Tim-3. At high concentrations of JNK inhibitor IL-2 production was decreased in both WT and CD28^A210P^ CD8^+^ T cells (**Fig 4H**). However, CD28^A210P^ production was not decreased to a level similar to WT as seen with MEK1/2 inhibitor, indicating although JNK activity promotes IL-2 production, JNK activity is not different between WT and CD28^A210P^ cells. Similar to results seen with MEK1/2 inhibitor, early IFNγ production is similar between WT and CD28^A210P^ T cells and, although high dose JNK inhibition can decrease IFNγ production in both WT and CD28^A210P^ T cells, this decrease is not significantly different between WT and CD28^A210P^ T cells (**Fig 4I**). In agreement with the pattern of inhibition of IL-2 production (**Fig 4H**), PD-1/Tim-3 are inhibited by JNK inhibition in both WT and CD28^A210P^ mice, but even at a high dose JNK inhibition PD-1^+^ Tim-3^+^ frequency remains higher CD28^A210P^ T cells compared to WT (**Fig 4J**), indicating JNK-dependent inhibitory receptor upregulation is not different between WT and CD28^A210P^ CD8^+^ T cells. Together these results suggest following αCD3/αCD28 activation, the CD28 A210P mutation increases MEK1/2-dependent induction of JunB, and this enhanced induction promotes IL-2 production and effector response, but also the upregulation of inhibitory receptors to limit the magnitude.

### Enhancing CD28 signaling does not affect numbers of effector/exhausted T cells during late chronic infection

CD28 signaling is required for effector/exhausted T cell differentiation during late chronic LCMV infection (*11*). We hypothesized enhancing CD28 C-terminal proline-rich domain signaling through the A210P mutation would promote promoting effector/exhausted T cell differentiation. We also wondered if PD-1 blockade would synergize with enhanced CD28 signaling to further promote effector/exhausted T cell output. To examine outcomes of enhanced CD28 signaling on CD8^+^ T cell exhaustion we utilized the CD4 depletion model of LCMV clone 13 infection with or without 2 weeks of PD-L1 blockade to reinvigorate the CD8^+^ T cell response (**Fig 5A**). At this late time point there is no difference in effector/exhausted CD8^+^ cell frequencies or absolute counts (**Fig 5B**) between WT and CD28^A210P^ mice. Surprisingly, although PD-L1 blockade increases the output of effector/exhausted cells as expected (**Fig 5B**), it does not synergize with enhanced CD28 signaling to further increase CD28^A210P^ effector/exhausted cell output (**Fig 5B**). Supporting a lack of synergy of enhanced CD28 signaling and PD-1 blockade, anti-PD-L1 increases the numbers of IFNγ-producing effector/exhausted cells in CD28^A210P^ mice to a number similar to WT mice (**Fig 5C**). Ratios of effector to exhausted cells are also similar between WT and CD28^A210P^ mice at this time point (**Fig 5D**). Furthermore, although PD-L1 blockade improves CD28^A210P^ viral control in the liver, viral burdens in peripheral organs are not different when comparing treatment matched WT and CD28^A210P^ mice (**Fig 5E**). These data, along with previous reports, suggest CD28 signaling is required for effector/exhausted differentiation in response to PD-L1 blockade during the chronic phase (*9, 10*), but intrinsically enhancing C-terminal CD28 signaling through the CD28 A210P mutation does not further promote effector/exhausted CD8^+^ T cell differentiation.

**Figure 5).**
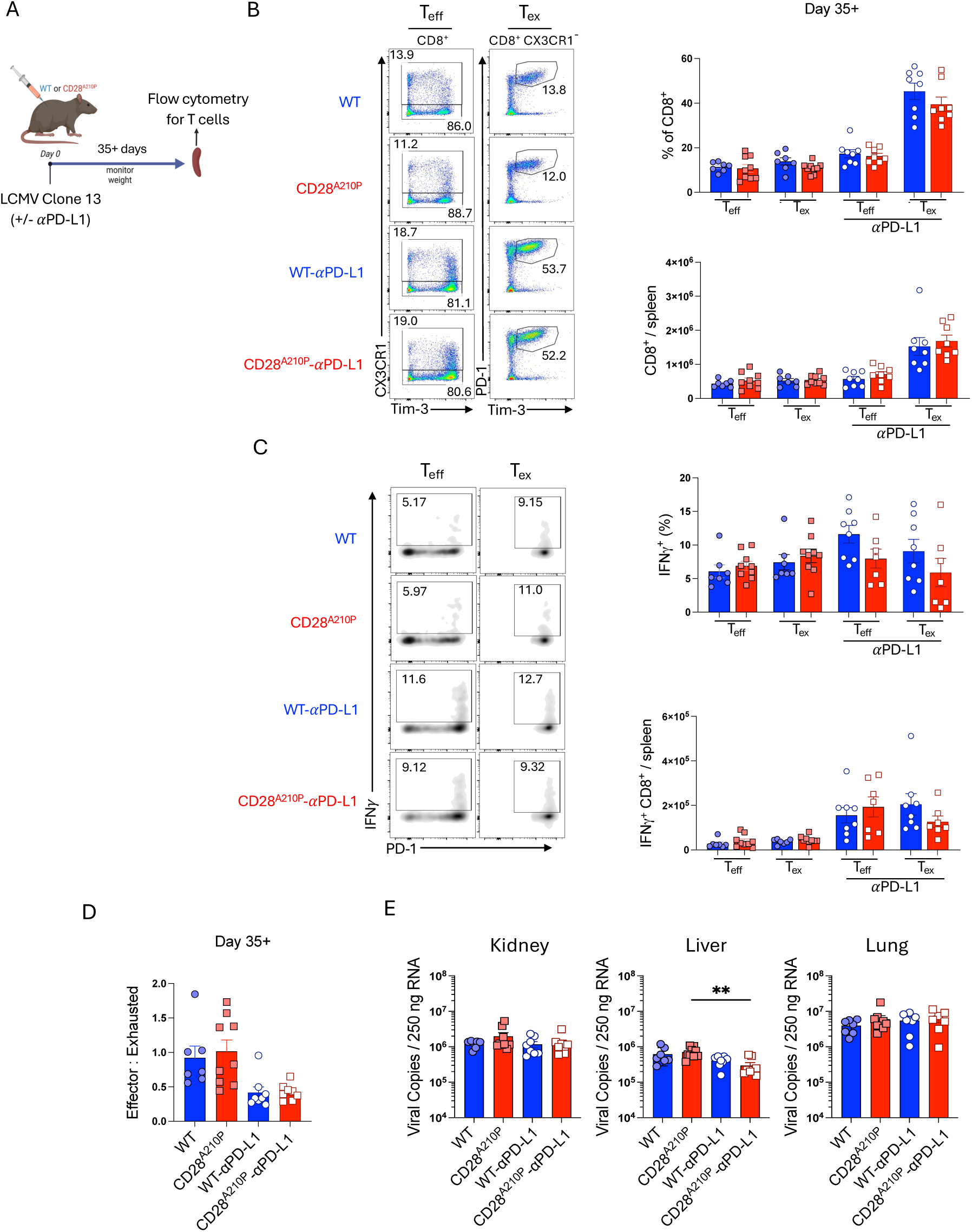
Enhancing CD28 signaling does not affect numbers of effector/exhausted T cells during late chronic infection. **A)** Experimental timeline of LCMV clone 13 infection with CD4-depletion. Anti-CD4 (GK1.5) was given I.P. (250 μg/mouse) on days –1 and +1. For experiments with anti-PD-L1 (10F.9G2) administration, 5 injections of anti-PD-L1 were administered I.P. (200 μg/injection) every 3 days for 2 weeks prior to analysis. **B)** Representative gating and quantification of frequencies and absolute counts of splenic CD8^+^ cell subsets on day 35+ of LCMV clone 13 infection. CD8^+^ subsets: effector (CX3CR1^+^) exhausted (CX3CR1^-^ PD-1^+^ Tim-3^-^). **C)** Representative gating and quantification of frequencies and absolute counts of IFNγ^+^ splenic CD8^+^ cell subsets on day 35+ of LCMV clone 13 infection. CD8^+^ subsets: effector (CX3CR1^+^) exhausted (CX3CR1^-^ PD-1^+^ Tim-3^-^). Splenocytes were re-stimulated ex vivo with LCMV GP33 peptide for 4 hours in the presence of GolgiPlug, followed by FACS staining. Data pooled from 2 independent experiments. **D)** Ratio of CD8^+^ effector (CX3CR1^+^) to exhausted (CX3CR1^-^ PD-1^+^ Tim-3^+^) splenocytes 35+ days post LCMV clone 13 infection. **E)** RT-qPCR quantification of viral copies in Kidney, liver, and lung tissue from LCMV clone 13 infected mice 35+ days post infection. Data are pooled from two independent experiments with 3-5 per group, dots show individual mice, significance assessed by Student’s t-test or one-way ANOVA, * = p < 0.05, ** = p < 0.01, *** = p < 0.001.

### Enhanced CD28 C-terminal signaling increases numbers and response to PD-L1 blockade during chronic infection

Stem-like Tpex CD8^+^ T cells arise during chronic infection and sustain the CD8^+^ effector response via asymmetric cell division, a process that is enhanced by PD-L1 blockade (*45*). 18 days post infection with chronic LCMV (**Fig 6A**), both WT and CD28^A210P^ mice have similar numbers of splenic Tpex (**Fig 6B**, S6A). Although total numbers of Tpex are similar at this time, CD28^A210P^ Tpex display enhanced IFNγ production upon ex vivo GP33 peptide stimulation (**Fig 6B**, S6B), suggesting enhanced CD28 signaling preserves some Tpex functionality at this time. However, CD28^A210P^ CD8^+^ T_eff_/T_ex_ ratio is decreased compared to WT cells (**Fig 3H**). We next asked how enhanced CD28 signaling would affect the numbers and functionality of Tpex late during chronic infection and if enhanced CD28 signaling in Tpex would synergize with PD-L1 blockade (**Fig 6C**). We confirmed that the CD28 A210P mutation does not alter CD28 surface expression among the various CD8^+^ T cell subsets involved in the chronic response phase of LCMV infection (**Fig 6D**). As previously reported (*11, 46*), CD28 surface expression is higher on Tpex relative to effector cells and exhausted cells. At day 35 post infection, CD28^A210P^ Tpex were maintained at a higher number than WT Tpex (**Fig 6E**, S6C), in agreement with the study by Humblin et al. that CD28 is required for the maintenance of Tpex during late chronic infection. As previously reported (*45*), PD-L1 blockade increased numbers of Tpex in WT mice relative to untreated controls. CD28^A210P^ Tpex were also increased in numbers relative to untreated, but after anti-PD-L1 treatment, CD28^A210P^ Tpex still outnumbered WT Tpex (**Fig 6E**, S6C), suggesting enhanced CD28 signaling further promotes increased numbers of Tpex with PD-L1 blockade. Without PD-L1 blockade, CD28^A210P^ Tpex produce more IFNγ following ex vivo GP33 peptide stimulation (**Fig 6E**, S6D). Upon PD-L1 blockade, IFNγ^+^ Tpex were increased in both WT and CD28^A210P^ splenocytes and there was a trend towards more IFNγ^+^ CD28^A210P^ Tpex but this greater increase did not reach statistical significance (**Fig 6E**, S6D). Overall enhancing CD28 signaling via the A210P mutation promotes increased numbers of Tpex during chronic LCMV infection and anti-PD-L1 treatment further increased numbers of Tpex (**Fig 6F**).

**Fig 6).**
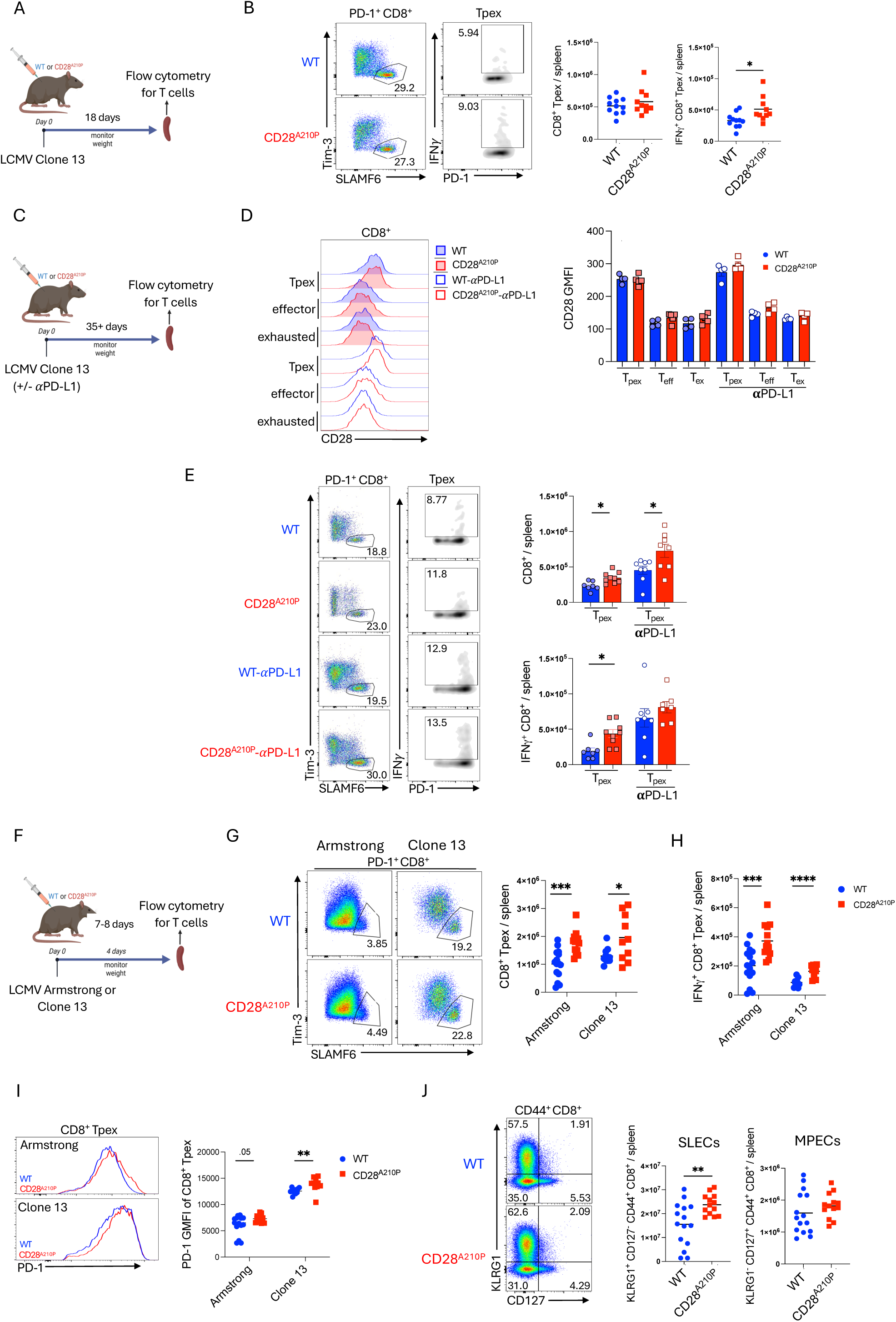
Enhancing CD28 increases Tpex differentiation and long-term maintenance without compromising memory precursors. **A)** Experimental timeline of LCMV Armstrong or clone 13 infection with CD4-depletion. Anti-CD4 (GK1.5) was given to indicated mice I.P. (250 μg/mouse) on days –1 and +1. **B)** Representative gating and quantification of absolute numbers of CD8^+^ Tpex (PD-1^+^ SLAMF6^+^ Tim-3^-^) 18 days post infection. For assessment of IFNγ potential, splenocytes were stimulated with LCMV GP33 peptide ex vivo for 4 hours in the presence of GolgiPlug. **C)** Experimental timeline of LCMV clone 13 infection with CD4-depletion. Anti-CD4 (GK1.5) was given I.P. (250 μg/mouse) on days –1 and +1. For mice treated with anti-PD-L1 (10F.9G2), 5 injections of anti-PD-L1 were administered I.P. (200 μg/injection) every 3 days for 2 weeks prior to analysis. **D)** Representative histograms and GMFIs of CD28 on CD8^+^ Tpex (PD-1^+^ SLAMF6^+^ Tim-3^-^), effectors (CX3CR1^+^), and exhausted (CX3CR1^-^ PD-1^+^ Tim-3^+^) cells 35+ days post LCMV infection. **E)** Representative gating and quantification of absolute counts of splenic CD8^+^ Tpex (PD-1^+^ SLAMF6^+^ Tim-3^-^), effectors (CX3CR1^+^), and exhausted (CX3CR1^-^ PD-1^+^ Tim-3^+^) cells 35+ days post LCMV infection. To determine absolute counts of IFNγ-producing CD8^+^ cells, 4-hour ex vivo LCMV GP33 peptide restimulation in the presence of GolgiPlug was performed. **F)** Quantification of absolute counts of splenic CD8^+^ Tpex (PD-1^+^ SLAMF6^+^ Tim-3^-^) 8-, 18-, or 42-days post infection with LCMV clone 13 with or without anti-PD-L1 treatment for two weeks. **G)** Experimental timeline of LCMV Armstrong or clone 13 infection. Anti-CD4 (GK1.5) was given to indicated mice I.P. (250 μg/mouse) on days –1 and +1. **H)** Representative gating and absolute counts of splenic CD8^+^ Tpex (PD-1^+^ SLAMF6^+^ Tim-3^-^) 7-8 days post infection. **I)** Absolute counts of IFNγ-producing CD8^+^ Tpex (PD-1^+^ SLAMF6^+^ Tim-3^-^) following 4-hour ex vivo LCMV GP33 peptide restimulation in the presence of GolgiPlug. **J)** Representative histograms and GMFIs of PD-1 on Tpex following LCMV infection. **K)** Representative gating and quantification of absolute numbers of CD8^+^ SLECs (CD44^+^ KLRG1^+^ CD127^-^) and MPECs (CD44^+^ KLRG1^-^ CD127^+^). Data are pooled or representative of at least two experiments with 3-6 per group, dots show individual mice, significance assessed by Student’s t-test or one-way ANOVA, * = p < 0.05, ** = p < 0.01, *** = p < 0.001.

### Enhanced CD28 C-terminal signaling increases early Tpex-like generation in acute and chronic infection without compromising memory precursors

Stem-like CD8^+^ T cells (Tpex) have mostly been implicated in conditions of chronic antigen stimulation (*11, 47*), but the presence of Tpex-like CD8 progenitors in acute infection was only recently described (*46, 48, 49*). As Tpex express high levels of *cd28* (*46*), we hypothesized early differentiation and function of Tpex may be affected by the enhanced CD28 signaling present in CD28^A210P^ mice. Following infection with LCMV Armstrong or Clone 13 (**Fig 6G**), CD28^A210P^ spleens had increased frequencies and numbers of Tpex-like CD8^+^ cells (PD-1^+^ SLAMF6^+^ Tim3^-^) (**Fig 6H**, S6E) compared to WT mice. Upon ex vivo stimulation with GP33 peptide CD28^A210P^ splenocytes contained more IFNγ^+^ and polyfunctional Tpex-like cells, suggesting functionality of the more abundant cells (**Fig 6I**, S6F-G). Like PD-1^+^ Tim-3^+^ CD8^+^ cells (**Fig 3C**), CD28^A210P^ Tpex display increased PD-1 surface expression compared to WT Tpex (**Fig 6J**), suggesting enhanced CD28 signaling also induces early cell-intrinsic regulation in Tpex.

Tpex-like cells arising during acute infection were shown to be a separate population from canonical short-lived effector cells (SLECs) and memory precursor effector cells (MPECs) (*48*), hence we asked if the increased differentiation of these cells in CD28^A210P^ mice affected the pool of SLECs or MPECs compared to WT mice. We examined SLECs (CD44^+^ KLRG1^+^ CD127^-^) and MPECs (CD44^+^ KLRG1^-^ CD127^+^) in WT and CD28^A10P^ mice 7 days post infection with LCMV Armstrong. In agreement with previous experiments (**Fig 2**), enhancing CD28 signaling increased SLEC differentiation (**Fig 6K**, S6H). Interestingly, frequencies of MPECs were decreased in CD28^A210P^ mice but total numbers of MPECs were similar to WT (**Fig 6K**, S6H). Together, these data suggest enhancing CD28 signaling during initial CD8^+^ T cell activation results in greater effector and stem-like T cell differentiation without perturbing early memory precursor induction.

## Discussion

Studies using deletion and targeted loss of function have led to consensus on the key CD28 signaling domains but sometimes conflicting conclusions of the function and relative contributions of these domains, often depending on the species and whether cell lines or primary cells are used (*4, 5*). Here we have shown that a species-specific difference in the amino acid flanking the C-terminal proline-rich domain PYAP profoundly alters the CD8 T cell response in mice, concurring with a prior study showing that alanine to proline substitution at the PYAP-proximal amino acid enhanced CD4 T cell response to CD28 autonomous signaling with a superagonist antibody (*30*). Primates including humans have an additional proline that enhances the function of the PYAP domain while mice, and indeed most mammals, have an inert alanine in that position which dulls CD28 function compared to human T cells in vitro (*30*). From an evolutionary point of view, it will be interesting to consider why primates have adopted this ‘stronger’ mode of CD28 signaling. A recent study found interspecies differences in strength of PD1 inhibition, with human PD1 being more inhibitory than mouse (*50*), further indicating that evolution of activating and inhibitory costimulatory receptor signaling that should be considered for pre-clinical studies targeting a specific signaling pathway.

CD28^A210P^ mice with the human PYAPP variant support the idea that enhanced inflammatory response to CD28 superagonist in humans is related to interspecies differences in the molecule itself. CD28^A210P^ mice did not develop life-threatening severity of cytokine storm experienced by humans, which we speculate could be due to the ultraclean environment in which our superagonist experiments were performed. ‘Wildling’ mice on the same C57Bl/6 background but with diverse microbial exposure demonstrated a remarkably similar reduction in Treg and increased inflammatory profile to CD28^A210P^ (*24*), suggesting that the combination of microbial-driven effector and memory T cells as well as enhanced CD28 signaling contributed to the unexpectedly catastrophic outcome of a CD28SA clinical trial in healthy volunteers. The unexpected observation of a striking increase in activated CD8 T cells in CD28^A210P^ compared to WT mice, suggests these cells could be an additional contributor that was underappreciated in preclinical mouse studies.

Enhancing CD28 function at the PYAP domain in CD28^A210P^ mice altered outcomes of physiological CD8 T cell stimulation with TCR and CD28 in vitro and during viral infection in vivo. Increased numbers of effector T cells was a somewhat predicted outcome of increasing CD28 strength. However, during chronic infection the increased effector response was rapidly outweighed by an acceleration of terminal T cell exhaustion in CD28^A210P^ mice, and the appearance of inhibitory receptors was accelerated even 24hr post-activation in vitro. T cell receptor affinity and cytokine signaling have previously been implicated as factors driving exhaustion in CD8 T cells during chronic stimulation (*51, 52*), our findings add CD28 co-stimulation strength as an additional regulator of exhaustion during the establishment of chronic infection.

If a virus is cleared from the host, the effector response rapidly subsides and a small population of circulating and tissue-resident memory T cells remain. Memory cells are not merely leftovers, in fact the fate of the memory pool is established remarkably early during the acute response (*46*). For CD8 T cells, strong glycolysis-driving signals have been shown to promote more effector cells at the expense of memory precursors (*53*). Hence, we predicted that the increased activation and effector cell generation might impair memory precursor formation in CD28^A210P^ T cells. This was not the case during acute infection with LCMV Armstrong, and instead it appears that increased CD8 T cell biomass early in the response allows increased effectors without impairing memory formation. The Akt-mTOR pathway is critical in driving enhanced glycolysis in T cells. We identified MEK1/2 and JunB as key contributors to both early activation and upregulation of inhibitory receptors in CD28^A210P^ T cells; in contrast the Akt pathway appeared unaffected corresponding to the N-terminal PI3K-binding domain being the key driver of PI3K and Akt phosphorylation (*5*). This fits with studies showing increased Vav1 recruitment by PYAPP (*30*), and that overexpression of Vav1 drives MEK-dependent JunB for increased IL-2 (*54*), while MEK1/2 inhibition limits JunB activity to prevent T cell exhaustion (*39, 41*). Individual signal transduction pathways and transcription factors do not operate in a vacuum during T cell activation, and so it is expected that additional TCR and cytokine-driven pathways intersect with and further enhance MEK1/2 driven enhancement of JunB downstream of PYAPP at later timepoints.

In contrast to effector and exhausted CD8 T cells, stem-like progenitor (Tpex) cells were increased at both early and late timepoints of LCMV cl13 infection in CD28^A210P^ mice, and were further increased in numbers by PDL-1 blockade. Tpex have been extensively studied under chronic conditions, but recent reports show that cells bearing Tpex characteristics arise separately from canonical memory precursors in the early stages of acute viral infection (*46, 48, 49*). Early stage Tpex-like cells have increased chromatin accessibility for AP-1 family members including JunB, relative to memory precursors (*48, 55*), perhaps explaining how CD28^A210P^ CD8 T cells are able to generate enhanced Tpex without affecting MPEC numbers. As antigen is cleared, Tpex adopt properties of Tcm cells (*46*), suggesting that CD28-mediated modulation of the Tpex pool could unequally alter the magnitude of subsequent memory T cell pools to enhance Tcm.

During chronic stimulation scenarios, Tpex act as a reservoir by undergoing asymmetric cell division to replenish cytotoxic effectors while also undergoing self-renewal (*47*). Tpex maintain notably high expression of CD28, and conditional deletion studies have shown that CD28 signaling is required to sustain Tpex during the chronic phase of LCMV clone 13 infection (*11*). CD28 signaling is thought to be the primary target ‘released’ by PD-L1 blockade to allow T cell proliferation during chronic antigen stimulation (*9, 10*). Furthermore, in vitro stimulation of Tpex with anti-CD28 increases effector output in a glycolysis-dependent manner after transfer into chronically infected animals (*11*). Accordingly, inhibiting mTOR favors early Tpex generation, and during PD-L1 blockade mTOR inhibition prevents Tpex to effector/exhausted differentiation (*56, 57*). Building on the idea that different CD28 signaling pathways drive different T cell fates, we found that enhancing CD28 activity via the PYAPP domain favors Tpex generation and renewal; however, it did not lead to increased effector T cells to any greater extent than PYAPA T cells following anti-PD-L1 treatment. This leads us to speculate that CD28 is necessary but may not be sufficient in humans (with enhanced PYAPP signal domain) to elicit strong effector generation from Tpex. Instead, these data support further targeting of Tpex by inclusion of signals further able to stimulate the Akt-glycolytic pathway for effector generation, for example through OX40 which is highly expressed on Tpex and enhances effector output when agonized (*58, 59*). Alternatively, one could test whether MEK1/2 inhibition along with anti-PD1 skews the effector to exhaustion ratio more favorably. Further, co-blockade of inhibitory molecules such as Lag-3 with PD-1 have been found to improve effector output during chronic LCMV (*60*), and it would be interesting to compare the effect of dual blockade on human CD28 signaling.

Another key therapeutic area where interspecies differences in CD28 signaling could be considered is in targeting of cancer, autoimmune, and infected cells for removal by T cells transfected with a chimeric antigen receptor (CAR). CAR-T receptors typically contain signaling domains of costimulatory molecules. In human CAR-T cells, incorporation of CD28 signaling domains increases inhibitory receptor expression compared to 4-1BB signaling (*61*), and silencing of the CD28 PI3K-Akt binding domain decreases exhaustion in vitro (*18*). However, in mouse CAR-T constructs incorporation of only the PYAP signaling domain from CD28 improved survival in vivo (*16*). With the caveat that CAR signaling is not directly comparable to TCR/CD28 stimulation, our data support the idea that modulating the balance of Akt and JunB activation via engineering of the CD28 and other co-stimulatory signaling domains may be a tool to balance activation/effector function with longevity for CAR-T efficacy.

In summary, this report introduces a new model to study CD28 signaling dynamics and a tool to investigate emerging T cell-based therapies for cancer, autoimmunity, and chronic infection that may be more relevant to humans. Just a single amino acid substitution adjacent to the PYAP domain to ‘humanize’ mouse CD28 signaling resulted in striking increases in CD8 T cell activation but also revealed roles for CD28 in regulating the balance of effector, exhausted, and progenitor populations that are set in place early during the response to viral infection. The outcomes of enhancing CD28 signaling in our murine studies support boosting PD1 immunotherapy by additional (non-CD28 dependent) targeting of Tpex that are particularly increased and sustained via enhanced PYAPP signaling.

## Materials and Methods

### Animals

C57Bl/6 mice were obtained from the Jackson Laboratory. CD28^A210P^ mice were generated via microinjection of C57BL/6L fertilized embryos with Cas9 protein, CD28 A210 sgRNA, and ssODN ultramer CD28-A210P-HDR. Embryos were transferred to the oviducts of pseudopregnant female recipients. Resulting offspring were genotyped to detect knock in, and positive mice were DNA sequenced to confirm insertion sequence and aligned with C57BL/6J genomic reference for identification of off-target alterations. Knock-in mice were back crossed with C57BL/6J for five generations. Both male and female mice were used for all experiments. Experiments utilized age and sex matched controls of 6–12-week-old mice. Mice were housed under specific pathogen–free conditions in an American Association of Laboratory Animal Care (AAALAC)-approved facility at the University of Pittsburgh and Cornell University. Protocols were approved by the University of Pittsburgh and Cornell university Institutional Animal Care and Use Committees and adhered to guidelines in the Guide for the Care and Use of Laboratory Animals of the National Institutes of Health.

### CD28 superagonist antibody

Anti-CD28 (D665) monoclonal antibody was purchased from BioXcell. For in vivo administration antibody was diluted in sterile PBS and 300 μg/mouse was administered via I.P. injection.

### In vitro T cell activation

Flat bottom tissue culture treated plates were coated with indicated concentration of agonistic monoclonal antibodies diluted in sterile PBS for T cell activation. Anti-CD3ε (145-2C11) and anti-CD28 (37.51) were purchased from BioXcell. Soluble cytokines (R&D biosystems) were added in indicated cultures. CD4^+^ or CD8^+^ T cells were purified from single cell suspensions of spleens and lymph nodes using magnetic isolation kits (Miltenyi) following manufacturer protocols.

### LCMV Armstrong infection

Virus aliquots were thawed and diluted in sterile RPMI. Mice were infected with 2×10^5 pfu via I.P. injection.

### LCMV Clone 13 infection

The Clone 13 strain of LCMV was propagated in baby hamster kidney (BHK-21) cells and supernatant harvested for storage at –80C. Virus titer was determined by fluorescent focus assays (FFA) on murine embryonic fibroblast (MEF) cells. Virus aliquots were thawed and diluted in sterile RPMI. Mice were infected with 2.6×10^6 pfu via retro-orbital (I.V.) injection. For indicated experiments CD4-depletion was performed via I.P. injection of 250 μg monoclonal anti-CD4 clone GK1.5 (BioXcell) on days –1 and +1. In indicated experiments 200 μg of anti-PD-L1 (10F.9G2) was administered via I.P. injections every third day for two weeks.

### LCMV viral quantification

LCMV clone 13 infected mice were euthanized via CO2 euthanasia and specified organs were snap frozen in liquid nitrogen for later processing. Tissues were later weighed and placed in trizol reagent before homogenization. RNA was then extracted using Qiagen RNeasy mini kit according following manufacturer protocol. 2 μg of RNA was converted to cDNA using High-Capacity cDNA Reverse Transcription Kit according to manufacturer protocol. cDNA was then diluted to 250 ng/μL for use in qPCR reaction. Quantabio SYBR Green RT-qPCR mix was used for reactions according to manufacturer protocol. Primers for LCMV GP F-CATTCACCTGGACTTTGTCAGACTC and GP R-GCAACTGCTGTGTTCCCGAAAC as described in (McCausland and Crotty, 2008) were purchased from IDT. Standard curves were generated using serial dilutions of a gene fragment (gblocks IDT) derived from Lymphocytic choriomeningitis virus clone 13 segment S, with the following sequence: 5’-AGA GAA GAC TAA GTT CCT CAC TAG GAG ACT AGC GGG CAC ATT CAC CTG GAC TTT GTC AGA CTC TTC AGG GGT GGA GAA TCC AGG TGG TTA TTG CCT GAC CAA ATG GAT GAT TCT TGC TGC AGA GCT TAA GTG TTT CGG GAA CAC AGC AGT TGC GAA ATG CAA TGT AAA TCA TGA TGA AGA ATT CTG TG 3’ as described in (Zander et al. 2022).

### ELISA

ELISA kits for IL-2 and IFNγ were purchased from Invitrogen and performed according to manufacturer protocol.

### Multiplex ELISA

Serum samples were shipped on dry ice and analyzed via multiplex ELISA (Eve Technologies).

### Western Blot and cytoplasmic/nuclear extraction

Cytoplasmic and Nuclear extracts were prepared according to manufacturer instruction manual (catalog#78833, Thermo Fisher) with protease inhibitors. Protein lysates were collected and mixed in sample buffer (1X), which were then incubated for 5 minutes at 95°C. SDS-PAGE was performed, and proteins were transferred to PVDF membranes, which were blocked with 0.1%TBS/Tween-20, 5% BSA for 1 hour at room temperature. The membrane was incubated with anti-Jun B (catalog# 3753S), anti-IRF-4 (catalog# 15106S), anti-BATF (catalog# 8638S), TBP (catalog# 77891S) Cell signaling Technology, overnight at 4°C. The blots were washed and then incubated with HRP-conjugated secondary antibody at room temperature for 1 hour. Membranes were developed with the Pierce ECL substrate kit (catalog# 32106 Thermo Fisher Scientific). Protein bands were visualized in Bio-Rad ChemiDoc Imaging System. Protein band intensity was quantified in Bio-Rad Image lab software.

### Flow cytometry

Single cell suspensions were obtained via tissue dissociation with 70 μm strainers followed by RBC lysis using ACK lysis buffer (Gibco). Viability staining was performed for 20 minutes at 4C using Ghost Dye UV450 (Tonbo Biosciences) in PBS. Surface staining was performed in FACS buffer (2% HI FBS, 5mM EDTA, PBS) for 30-60 minutes at 4C. Cells were then fixed with IC fixation buffer (Invitrogen) or FOXP3 Fixation buffer (Invitrogen) for 20 minutes at room temperature. For intracellular/nuclear staining cells were stained in perm/wash buffer (Invitrogen) for 45 minutes at 4C. For intracellular cytokine staining splenocytes were stimulated ex vivo in 96-well plates at 37C for 4 hours with PMA/Ionomycin or indicated peptides in the presence of Golgiplug (BD). For phospho-flow experiments cells were fixed in 4% PFA at 37C for 15 minutes and permeabilized with ice cold methanol on ice for at least 1 hour prior to staining. Stained cells were acquired on a BD FACSymphony A5 SE and analyzed using FlowJo software (BD).

### Flow Cytometry Reagents

Tetramers were obtained from the NIH tetramer Core (Emory). Flow cytometry antibodies were obtained from BD, Biolegend, Thermofisher, and Cell Signaling Technologies. The following clones were utilized: CD4 (RM4-5), CD44 (IM7), CD62L (MEL-14), CD8 (53-6.7), Ly108 (13G3), Tim-3 (RMT3-23), PD-1 (29F.1A12), ICOS(7E.17G9), KLRG1(2f1), CD127(SB/199), CD28(37.51), TCR-β(H57-597), CD69(H1.2F3), CD25(PC61), Helios(22F6), NRP1(3E12), CX3CR1(SA011F11), IFNγ (XMG1.2), TNF (MP6-XT22), IL-2 (JES6-5H4), Granzyme B (QA16A02), B220 (RA3-6B2), FOXP3 (FJK-16s), Ki67 (SolA15), CTLA-4 (UC10-4B9), CD45.1 (A20), CD45.2 (104), Nur77 (12.14), p-Akt (M89-61), p-Erk1/2 (197G2), p-S6 (D57.2.2E).

### Statistics

Experimental results were analyzed for significance using one-way analysis of variance (ANOVA) with Tukey’s multiple comparisons test (for multiple groups) or unpaired Student’s t test. Statistical analyses were performed using GraphPad Prism. P values are shown as *P < 0.05, **P < 0.01, ***P < 0.001, and ****P < 0.0001 where statistical significance was found, and all data are represented as means ± SEM. Individual points in graphs represent biological replicates (i.e., individual mice) pooled from multiple experiments unless otherwise indicated in figure legends.

## List of Supplementary Materials

Fig S1) Enhanced CD28 signaling in CD28A210P mice does not alter T cell development or homeostatic peripheral T cell populations

Fig S2) CD28^A210P^ mice undergo a proinflammatory response to CD28 superagonist antibody extended

Fig S3) Enhancing CD28 signaling increases the initial effector CD8^+^ T cell response to acute and chronic infection extended

Fig S4) Enhancing CD28 C-terminal signaling induces early upregulation of CD8^+^ T cell inhibitory receptors and expedites exhaustion extended

Fig S5) CD28^A210P^ T cells display enhanced JunB nuclear localization compared to WT while other CD28-dependent signaling is similar to WT T cells

Fig S6) Enhancing CD28 increases Tpex differentiation and long-term maintenance without compromising memory precursor extended

## Acknowledgments

Funding: NIH AI170773, AI144229, Cornell College of Veterinary Medicine. Services: We thank the staff of University of Pittsburgh Division of Laboratory Animal Research (DLAR) and Cornell University Center for Animal Resources and Education (CARE) for assistance with mouse husbandry and care. We acknowledge Sage Golden, Lamisa Tasneem and Natasha Zarrin for assistance with genotyping, and Summer LaPointe for manuscript editing. We would like to acknowledge the contributions of Chunming Bi and Zhaohui Kou in the Mouse Embryo Services Core, University of Pittsburgh, Department of Immunology for microinjection of zygotes to produce the CD28^A210P^ mice. We acknowledge the Cornell Institute of Biotechnology flow cytometry services and University of Pittsburgh Unified Flow Core for assistance with flow cytometry.

## FIGURE LEGENDS

**Fig S1).**
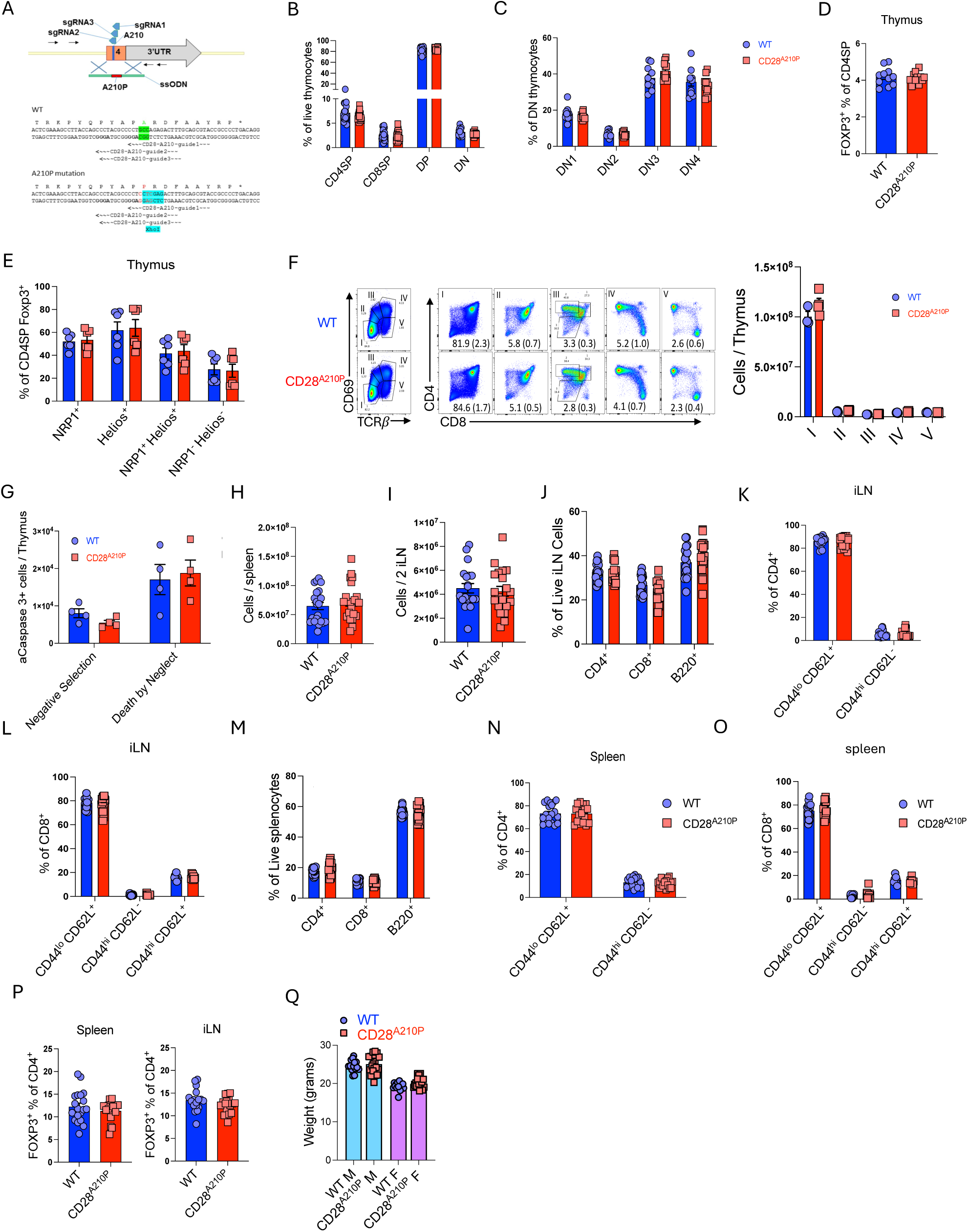
Enhanced CD28 signaling in CD28A210P mice does not alter T cell development or homeostatic peripheral T cell populations. **A)** Target DNA sequence of CRISPR/Cas9 sgRNAs. In blue: sequence for XhoI enzyme digestion to facilitate genotyping of CD28A210P mice. **B-G)** Flow cytometry analysis of adult thymocytes. **B)** Frequencies of thymocyte subpopulations **C)** Frequencies of DN subpopulations among the DN cells, Gated using CD44 and CD25. **D)** Frequencies of Foxp3^+^ among CD4SP thymocytes. **E)** Frequencies of NRP1^+^ and Helios^+^ cells among thymic Foxp3^+^ Tregs. **F)** Representative gating and absolute counts of thymic developmental subsets. **G)** Positively and negatively selected cells based on cleaved caspase-3 staining in signaled and not signaled thymocytes determined by CD5 and TCR-β. **H-P)** Flow cytometry analysis of adult spleens and inguinal lymph nodes. **H)** Absolute counts of splenocytes. **I)** Absolute counts of inguinal lymph node cells. **J)** Frequencies of CD4^+^, CD8^+^, and B220^+^ cells among live inguinal lymph node cells. **K)** Frequencies of CD44^hi^ CD62L^-^ and CD44^lo^ CD62L^+^ CD4^+^ cells in inguinal lymph nodes. **L)** Frequencies of CD44^lo^ CD62L^+^, CD44^hi^ CD62L^-^, and CD44^hi^ CD62L^+^ CD8^+^ splenocytes. **M)** Frequencies of CD44^lo^ CD62L^+^, CD44^hi^ CD62L^-^, and CD44^hi^ CD62L^+^ CD8^+^ cells in inguinal lymph nodes. **N)** Frequencies of CD4^+^, CD8^+^, and B220^+^ cells among live splenocytes. **O)** Frequencies of CD44^hi^ CD62L^-^ and CD44^lo^ CD62L^+^ CD4^+^ splenocytes. **P)** Frequencies of Foxp3^+^ of CD4^+^ cells in spleens and inguinal lymph nodes. **Q)** Body weights of adult male and female WT and CD28^A210P^ mice. Data are pooled or representative of at least two experiments with significance assessed by Student’s t-test or one-way ANOVA, * = p < 0.05, ** = p < 0.01, *** = p < 0.001.

**Fig S2).**
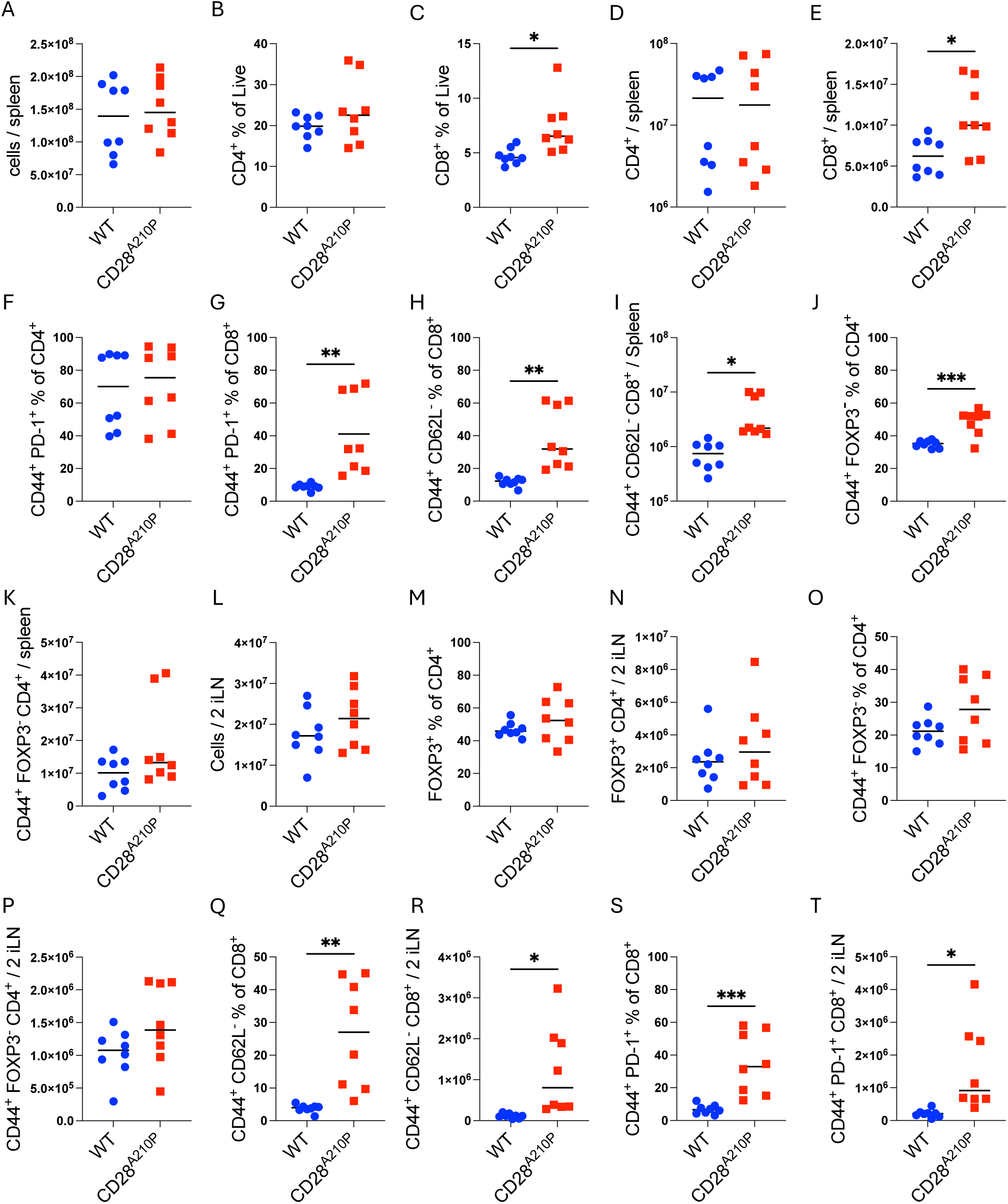
CD28^A210P^ mice undergo a proinflammatory response to CD28 superagonist antibody extended. **A-K**) Flow cytometric analysis of splenocytes from WT and CD28A210P mice 4 days post injection with CD28 superagonist antibody. Frequencies shown as % of parent population and absolute numbers calculated from total tissue cell counts. **L-T)** Flow cytometric analysis of cell from pooled 2 inguinal lymph nodes of WT and CD28A210P mice 4 days post injection with CD28 superagonist antibody. Frequencies shown as % of parent population and absolute numbers calculated from total tissue cell counts. Data are pooled from two independent experiments with significance assessed by Student’s t-test, * = p < 0.05, ** = p < 0.01, *** = p < 0.001.

**Fig S3).**
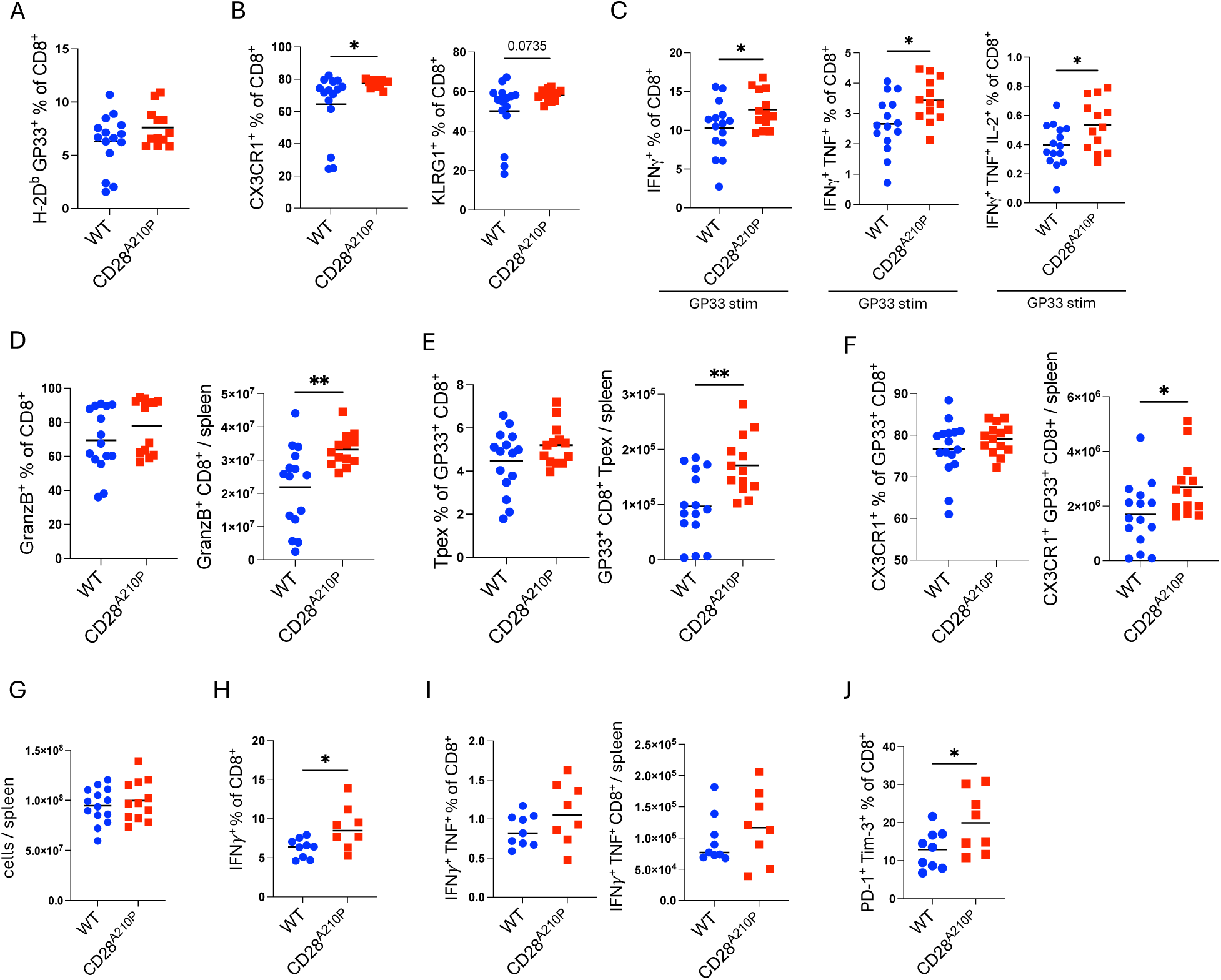
Enhancing CD28 signaling increases the initial effector CD8^+^ T cell response to acute and chronic infection extended. **A-F**) Flow cytometry data from WT and CD28A210P splenocytes 7 days post infection with LCMV Armstrong. For intracellular cytokine staining (C,D) splenocytes were restimulated ex vivo for 4 hours with GP33 peptide in the presence of GolgiPlug. GP33 tetramer staining were done for 1 hour at 4C. Tpex = CD8^+^ PD-1^+^ SLAMF6^+^ Tim-3^-^. GP33^+^ = H-2D^b^ GP33 tetramer^+^. **G-J)** Flow cytometry data from WT and CD28^A210P^ splenocytes 8 days post infection with LCMV clone 13. For intracellular cytokine staining (H,I) splenocytes were restimulated ex vivo for 4 hours with GP33 peptide in the presence of GolgiPlug. J) PD-1^+^ Tim-3^+^ frequencies among CD8^+^ cells. Data are pooled from or representative of two independent experiments with significance assessed by Student’s t-test, * = p < 0.05, ** = p < 0.01, *** = p < 0.001.

**Fig S4).**
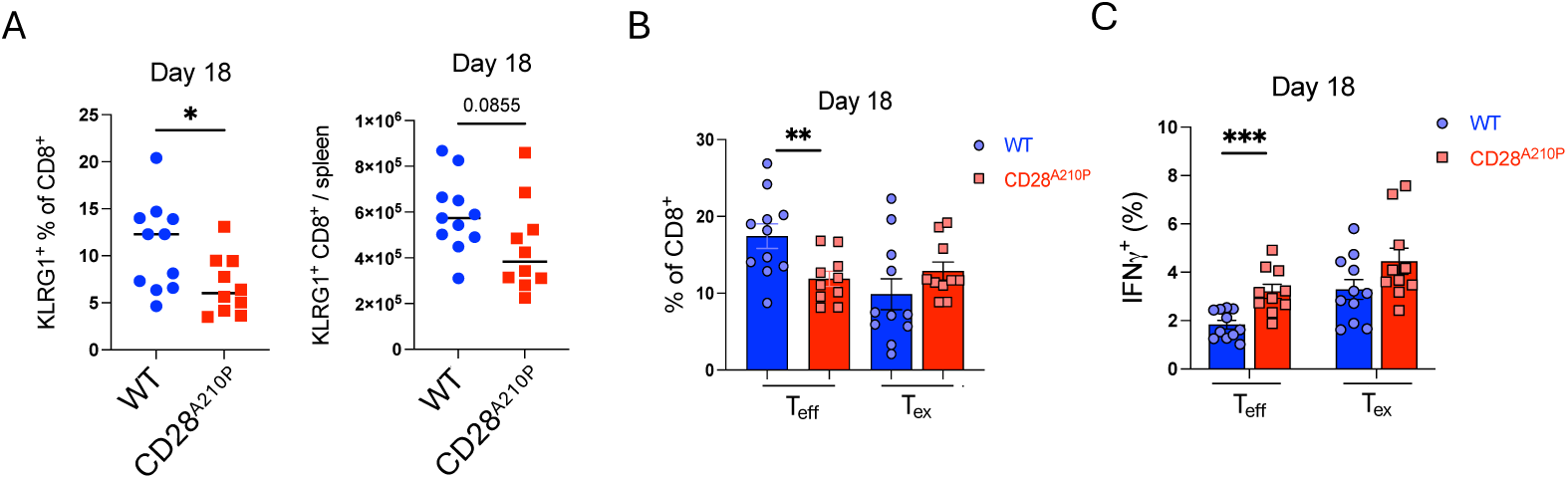
Enhancing CD28 C-terminal signaling induces early upregulation of CD8^+^ T cell inhibitory receptors and expedites exhaustion extended. **A-C**) Flow cytometry analysis of spleens from WT and CD28^A210P^ mice infected 18 days prior with LCMV clone 13. **A)** Frequencies and absolute numbers of KLRG1^+^ CD8^+^ cells. **B)** Frequencies of CD8^+^ effector and exhausted cells. **C)** Analysis of intracellular cytokine production of D18 LCMV clone 13 infected WT and CD28^A210P^ splenocytes restimulated ex vivo with GP33 peptide for 4 hours in the presence of GolgiPlug. Effector = CD8^+^ CX3CR1^+^ Exhausted = CD8^+^ CX3CR1^-^ PD-1^+^ Tim-3^+^. Data are pooled from two independent experiments with significance assessed by Student’s t-test, * = p < 0.05, ** = p < 0.01, *** = p < 0.001.

**Fig S5).**
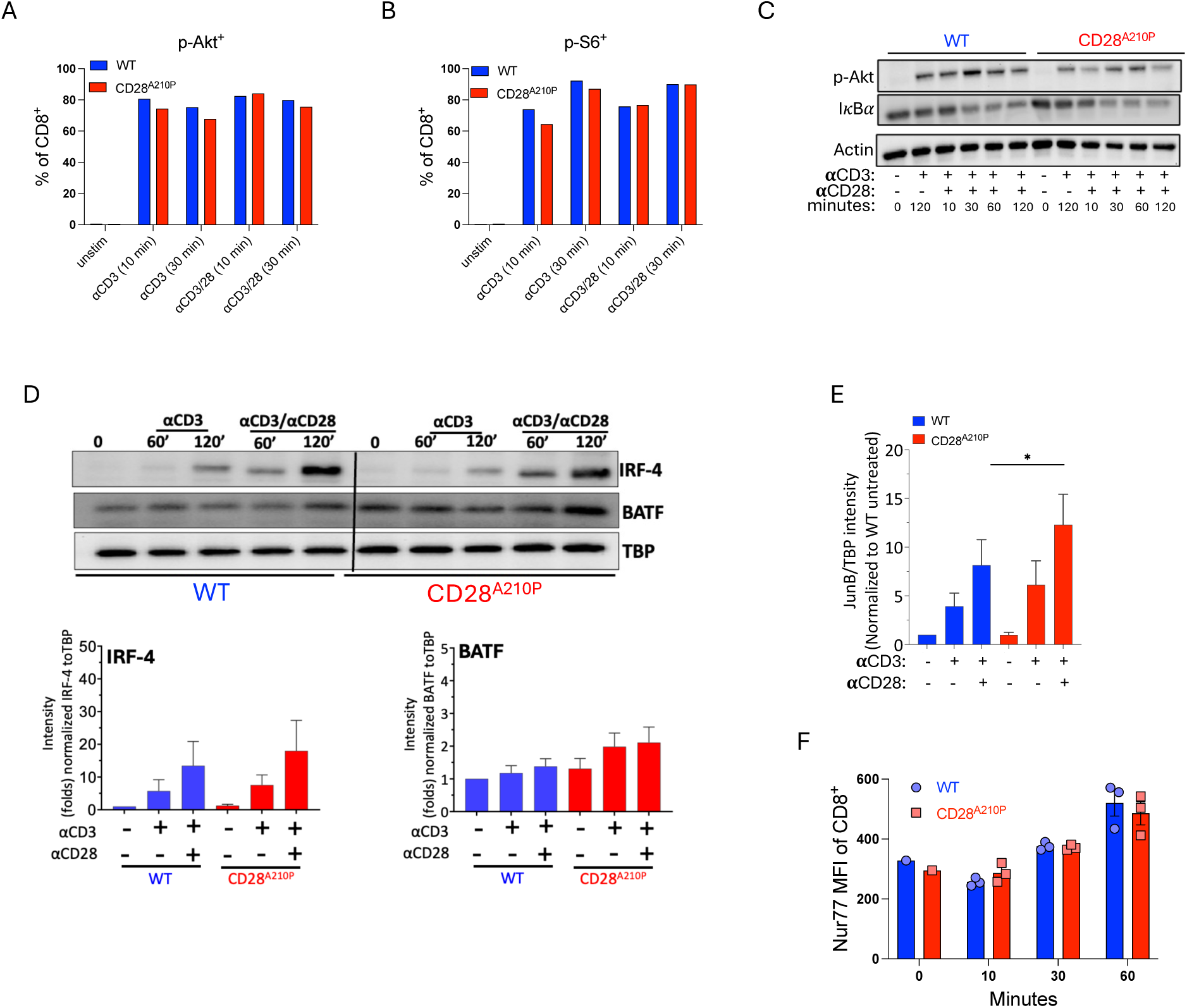
CD28^A210P^ T cells display enhanced JunB nuclear localization compared to WT while other CD28-dependent signaling is similar to WT T cells. **A-B**) WT or CD28^A210P^ CD8^+^ T cells were enriched from secondary lymphoid organs using MACS enrichment. Cells were then stimulated with plate bound agonistic anti-CD3/28 (1 μg/mL). Cells were stimulated for 10-30 minutes as indicated and then harvested for p-flow cytometric analysis of p-Akt and p-S6. Data representative of 3 independent experiments. **C-D)** WT or CD28^A210P^ CD8^+^ T cells were stimulated with plate bound agonistic anti-CD3/28 (5 μg/mL) as indicated. Cells were stimulated for up to 2 hours prior to lysis for immunoblot analysis. Akt phosphorylation (S473) and IKBα degradation assessed from cytoplasmic fractions and IRF4 and BATF assessed from nuclear extracts. C) is representative of 2 independent experiments and D) is representative of 3-4 independent experiments. **E)** WT or CD28^A210P^ CD8^+^ T cells were stimulated with plate bound agonistic anti-CD3/28 (5 μg/mL) as indicated. Cells were stimulated for 2 hours prior to nuclear extraction for immunoblot analysis, quantification of 5 pooled experiments shown. **F)** WT or CD28^A210P^ CD8^+^ T cells were stimulated with plate bound agonistic anti-CD3/28 (5 μg/mL) for indicated times. Nur77 staining among CD8^+^ cells was quantified. Data are pooled or representative of 3-4 independent experiments with significance assessed by Student’s t-test or one-way ANOVA, * = p < 0.05, ** = p < 0.01, *** = p < 0.001.

**Fig S6).**
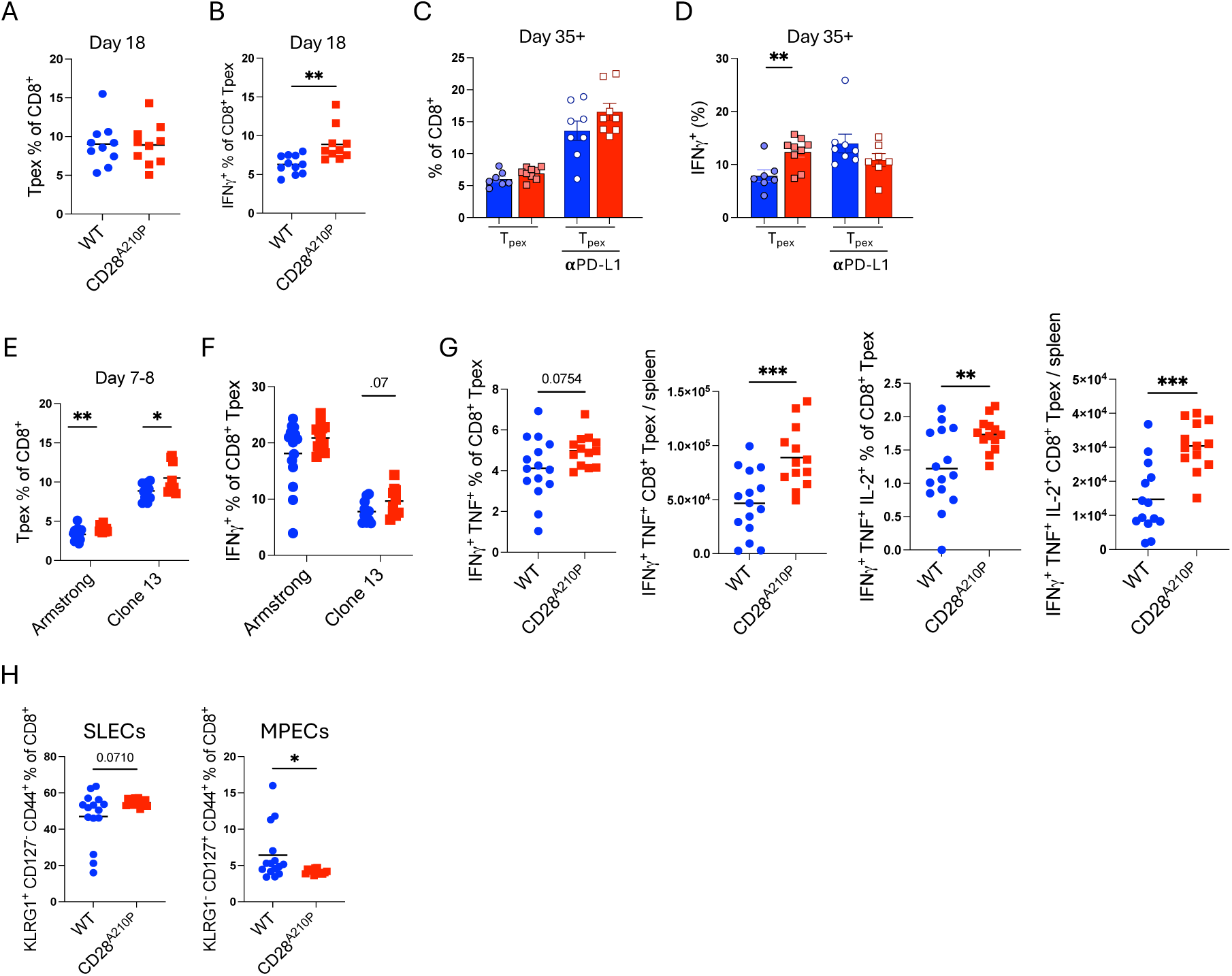
Enhancing CD28 increases Tpex differentiation and long-term maintenance without compromising memory precursor extended. **A-B**) WT or CD28^A210P^ mice were infected with LCMV Clone 13. 18 days post infection splenocytes were stained for flow cytometric analysis. **A)** Frequencies of CD8^+^ Tpex. **B)** Intracellular cytokine staining of splenocytes restimulated with LCMV GP33 peptide ex vivo for 4 hours in the presence of GolgiPlug. **C-D)** WT or CD28^A210P^ mice were infected with LCMV Clone 13 and 35+ days post infection splenocytes were stained for flow cytometric analysis. For mice treated with anti-PD-L1 (10F.9G2), 5 injections of anti-PD-L1 were administered I.P. (200 μg/injection) every 3 days for 2 weeks prior to analysis. **C)** Frequencies of Tpex among CD8^+^ cells. **D)** Frequencies of IFNγ^+^ cells following ex vivo GP33 peptide restimulation. **E-F)** Flow cytometric analysis of spleens from LCMV Armstrong or clone 13 infected WT and CD28^A210P^ mice (D7-8). **E)** Frequencies of Tpex among CD8^+^ cells. **F)** Frequencies of IFNγ^+^ cells following ex vivo GP33 peptide restimulation. **G-H)** Flow cytometry of spleens from LCMV Armstrong infected WT and CD28^A210P^ mice (Day 7). **G)** Frequencies and absolute counts of cytokine producing CD8^+^ Tpex following ex vivo GP33 peptide restimulation. **H)** Frequencies of SLECs and MPECs among CD8^+^ cells. For Tpex = CD8^+^ PD-1^+^ SLAMF6^+^ Tim-3^-^. Data are pooled from 2 independent experiments with significance assessed by Student’s t-test, * = p < 0.05, ** = p < 0.01, *** = p < 0.001.

## References

1. J. S. Burr et al., Cutting edge: distinct motifs within CD28 regulate T cell proliferation and induction of Bcl-XL. J Immunol 166, 5331–5335 (2001).

2. L. F. Dodson et al., Targeted knock-in mice expressing mutations of CD28 reveal an essential pathway for costimulation. Mol Cell Biol 29, 3710–3721 (2009).

3. P. G. Andres et al., Distinct regions in the CD28 cytoplasmic domain are required for T helper type 2 differentiation. Nat Immunol 5, 435–442 (2004).

4. J. H. Esensten, Y. A. Helou, G. Chopra, A. Weiss, J. A. Bluestone, CD28 Costimulation: From Mechanism to Therapy. Immunity 44, 973–988 (2016).

5. J. S. Boomer, J. M. Green, An enigmatic tail of CD28 signaling. Cold Spring Harb Perspect Biol 2, a002436 (2010).

6. Y. C. Cai et al., Selective CD28pYMNM mutations implicate phosphatidylinositol 3-kinase in CD86-CD28-mediated costimulation. Immunity 3, 417–426 (1995).

7. M. Watanabe, Y. Lu, M. Breen, R. J. Hodes, B7-CD28 co-stimulation modulates central tolerance via thymic clonal deletion and Treg generation through distinct mechanisms. Nat Commun 11, 6264 (2020).

8. C. W. Lio, L. F. Dodson, C. M. Deppong, C. S. Hsieh, J. M. Green, CD28 facilitates the generation of Foxp3(-) cytokine responsive regulatory T cell precursors. J Immunol 184, 6007–6013 (2010).

9. A. O. Kamphorst et al., Rescue of exhausted CD8 T cells by PD-1-targeted therapies is CD28-dependent. Science 355, 1423–1427 (2017).

10. E. Hui et al., T cell costimulatory receptor CD28 is a primary target for PD-1-mediated inhibition. Science 355, 1428–1433 (2017).

11. E. Humblin, et al., Sustained CD28 costimulation is required for self-renewal and differentiation of TCF-1(+) PD-1(+) CD8 T cells. Sci Immunol 8, eadg0878 (2023).

12. N. Prokhnevska et al., CD8(+) T cell activation in cancer comprises an initial activation phase in lymph nodes followed by effector differentiation within the tumor. Immunity 56, 107–124 e105 (2023).

13. F. Vincenti et al., Five-year safety and efficacy of belatacept in renal transplantation. J Am Soc Nephrol 21, 1587–1596 (2010).

14. N. Poirier et al., FR104, an antagonist anti-CD28 monovalent fab’ antibody, prevents alloimmunization and allows calcineurin inhibitor minimization in nonhuman primate renal allograft. Am J Transplant 15, 88–100 (2015).

15. K. P. Burke, A. Chaudhri, G. J. Freeman, A. H. Sharpe, The B7:CD28 family and friends: Unraveling coinhibitory interactions. Immunity 57, 223–244 (2024).

16. J. C. Boucher et al., CD28 Costimulatory Domain-Targeted Mutations Enhance Chimeric Antigen Receptor T-cell Function. Cancer Immunol Res 9, 62–74 (2021).

17. M. M. Honikel, S. H. Olejniczak, Co-Stimulatory Receptor Signaling in CAR-T Cells. Biomolecules 12, (2022).

18. S. Guedan et al., Single residue in CD28-costimulated CAR-T cells limits long-term persistence and antitumor durability. J Clin Invest 130, 3087–3097 (2020).

19. C. H. Lin, T. Hunig, Efficient expansion of regulatory T cells in vitro and in vivo with a CD28 superagonist. Eur J Immunol 33, 626–638 (2003).

20. M. Rodriguez-Palmero et al., Effective treatment of adjuvant arthritis with a stimulatory CD28-specific monoclonal antibody. J Rheumatol 33, 110–118 (2006).

21. N. Beyersdorf et al., Selective targeting of regulatory T cells with CD28 superagonists allows effective therapy of experimental autoimmune encephalomyelitis. J Exp Med 202, 445–455 (2005).

22. G. Suntharalingam et al., Cytokine storm in a phase 1 trial of the anti-CD28 monoclonal antibody TGN1412. N Engl J Med 355, 1018–1028 (2006).

23. L. K. Beura et al., Normalizing the environment recapitulates adult human immune traits in laboratory mice. Nature 532, 512–516 (2016).

24. S. P. Rosshart et al., Laboratory mice born to wild mice have natural microbiota and model human immune responses. Science 365, (2019).

25. N. Panoskaltsis et al., Immune reconstitution and clinical recovery following anti-CD28 antibody (TGN1412)-induced cytokine storm. Cancer Immunol Immunother 70, 1127–1142 (2021).

26. D. Eastwood et al., Monoclonal antibody TGN1412 trial failure explained by species differences in CD28 expression on CD4+ effector memory T-cells. Br J Pharmacol 161, 512–526 (2010).

27. N. E. McCarthy et al., Patients with gastrointestinal irritability after TGN1412-induced cytokine storm displayed selective expansion of gut-homing alphabeta and gammadeltaT cells. Cancer Immunol Immunother 70, 1143–1153 (2021).

28. L. D. Friend et al., A dose-dependent requirement for the proline motif of CD28 in cellular and humoral immunity revealed by a targeted knockin mutant. J Exp Med 203, 2121–2133 (2006).

29. J. S. Boomer, C. M. Deppong, D. D. Shah, T. L. Bricker, J. M. Green, Cutting edge: A double-mutant knockin of the CD28 YMNM and PYAP motifs reveals a critical role for the YMNM motif in regulation of T cell proliferation and Bcl-xL expression. J Immunol 192, 3465–3469 (2014).

30. N. Porciello et al., A non-conserved amino acid variant regulates differential signalling between human and mouse CD28. Nat Commun 9, 1080 (2018).

31. D. C. Fajgenbaum, C. H. June, Cytokine Storm. N Engl J Med 383, 2255–2273 (2020).

32. H. Frebel et al., Programmed death 1 protects from fatal circulatory failure during systemic virus infection of mice. J Exp Med 209, 2485–2499 (2012).

33. P. M. Odorizzi, K. E. Pauken, M. A. Paley, A. Sharpe, E. J. Wherry, Genetic absence of PD-1 promotes accumulation of terminally differentiated exhausted CD8+ T cells. J Exp Med 212, 1125–1137 (2015).

34. V. Atsaves, V. Leventaki, G. Z. Rassidakis, F. X. Claret, AP-1 Transcription Factors as Regulators of Immune Responses in Cancer. Cancers (Basel) 11, (2019).

35. A. Mondino et al., Defective transcription of the IL-2 gene is associated with impaired expression of c-Fos, FosB, and JunB in anergic T helper 1 cells. J Immunol 157, 2048–2057 (1996).

36. M. Yukawa et al., AP-1 activity induced by co-stimulation is required for chromatin opening during T cell activation. J Exp Med 217, (2020).

37. S. V. Janardhan, K. Praveen, R. Marks, T. F. Gajewski, Evidence implicating the Ras pathway in multiple CD28 costimulatory functions in CD4+ T cells. PLoS One 6, e24931 (2011).

38. P. J. R. Ebert et al., MAP Kinase Inhibition Promotes T Cell and Anti-tumor Activity in Combination with PD-L1 Checkpoint Blockade. Immunity 44, 609–621 (2016).

39. X. Wang et al., MEK inhibition prevents CAR-T cell exhaustion and differentiation via downregulation of c-Fos and JunB. Signal Transduct Target Ther 9, 293 (2024).

40. K. L. McGuire, M. Iacobelli, Involvement of Rel, Fos, and Jun proteins in binding activity to the IL-2 promoter CD28 response element/AP-1 sequence in human T cells. J Immunol 159, 1319–1327 (1997).

41. V. Verma et al., MEK inhibition reprograms CD8(+) T lymphocytes into memory stem cells with potent antitumor effects. Nat Immunol 22, 53–66 (2021).

42. G. Xiao, A. Deng, H. Liu, G. Ge, X. Liu, Activator protein 1 suppresses antitumor T-cell function via the induction of programmed death 1. Proc Natl Acad Sci U S A 109, 15419–15424 (2012).

43. S. J. Yun et al., The regulation of TIM-3 transcription in T cells involves c-Jun binding but not CpG methylation at the TIM-3 promoter. Mol Immunol 75, 60–68 (2016).

44. W. Li et al., CD28 signaling augments Elk-1-dependent transcription at the c-fos gene during antigen stimulation. J Immunol 167, 827–835 (2001).

45. A. L. Gill, et al., PD-1 blockade increases the self-renewal of stem-like CD8 T cells to compensate for their accelerated differentiation into effectors. Sci Immunol 8, eadg0539 (2023).

46. D. T. McManus et al., An early precursor CD8 T cell that adapts to acute or chronic viral infection. Nature, (2025).

47. D. Zehn, R. Thimme, E. Lugli, G. P. de Almeida, A. Oxenius, ’Stem-like’ precursors are the fount to sustain persistent CD8(+) T cell responses. Nat Immunol 23, 836–847 (2022).

48. T. Chu et al., Precursors of exhausted T cells are preemptively formed in acute infection. Nature, (2025).

49. E. Fagerberg et al., KLF2 maintains lineage fidelity and suppresses CD8 T cell exhaustion during acute LCMV infection. Science 387, eadn2337 (2025).

50. T. Masubuchi et al., Functional differences between rodent and human PD-1 linked to evolutionary divergence. Sci Immunol 10, eads6295 (2025).

51. E. J. Wherry, M. Kurachi, Molecular and cellular insights into T cell exhaustion. Nat Rev Immunol 15, 486–499 (2015).

52. A. Baessler, D. A. A. Vignali, T Cell Exhaustion. Annu Rev Immunol 42, 179–206 (2024).

53. M. D. Buck, D. O’Sullivan, E. L. Pearce, T cell metabolism drives immunity. J Exp Med 212, 1345–1360 (2015).

54. C. Charvet, P. Auberger, S. Tartare-Deckert, A. Bernard, M. Deckert, Vav1 couples T cell receptor to serum response factor-dependent transcription via a MEK-dependent pathway. J Biol Chem 277, 15376–15384 (2002).

55. B. Daniel et al., Divergent clonal differentiation trajectories of T cell exhaustion. Nat Immunol 23, 1614–1627 (2022).

56. K. A. Frauwirth et al., The CD28 signaling pathway regulates glucose metabolism. Immunity 16, 769–777 (2002).

57. S. Ando et al., mTOR regulates T cell exhaustion and PD-1-targeted immunotherapy response during chronic viral infection. J Clin Invest 133, (2023).

58. T. C. van der Sluis et al., OX40 agonism enhances PD-L1 checkpoint blockade by shifting the cytotoxic T cell differentiation spectrum. Cell Rep Med 4, 100939 (2023).

59. T. W. Kim et al., First-In-Human Phase I Study of the OX40 Agonist MOXR0916 in Patients with Advanced Solid Tumors. Clin Cancer Res 28, 3452–3463 (2022).

60. S. F. Ngiow et al., LAG-3 sustains TOX expression and regulates the CD94/NKG2-Qa-1b axis to govern exhausted CD8 T cell NK receptor expression and cytotoxicity. Cell 187, 4336–4354 e4319 (2024).

61. M. E. Selli et al., Costimulatory domains direct distinct fates of CAR-driven T-cell dysfunction. Blood 141, 3153–3165 (2023).

62. C. UniProt, UniProt: the Universal Protein Knowledgebase in 2025. Nucleic Acids Res 53, D609–D617 (2025).

